# Distinct and interdependent functions of three RING proteins regulate recombination during mammalian meiosis

**DOI:** 10.1101/2023.11.07.566091

**Authors:** Masaru Ito, Yan Yun, Dhananjaya S. Kulkarni, Sunkyung Lee, Sumit Sandhu, Briana Nuñez, Linya Hu, Kevin Lee, Nelly Lim, Rachel M. Hirota, Rowan Prendergast, Cynthia Huang, Ivy Huang, Neil Hunter

## Abstract

During meiosis, each pair of homologous chromosomes becomes connected by at least one crossover, as required for accurate segregation at the first division, and adjacent crossovers are widely separated thereby limiting total numbers. In coarsening models, crossover patterning results from nascent recombination sites competing to accrue a limiting pro-crossover RING-domain protein (COR) that diffuses between synapsed chromosomes. Here we delineate the localization dynamics of three mammalian CORs in mouse and determine their interdependencies. RNF212, HEI10 and a new member RNF212B show divergent spatiotemporal dynamics along synapsed chromosomes, including profound differences in spermatocytes and oocytes, that are not easily reconciled by elementary coarsening models. Contrasting mutant phenotypes and genetic requirements indicate that RNF212B, RNF212, and HEI10 play distinct but interdependent functions in regulating meiotic recombination and coordinating the events of meiotic prophase-I by integrating signals from DNA breaks, homolog synapsis, the cell-cycle, and incipient crossover sites.

**SIGNIFICANCE:** Meiosis produces haploid gametes by precisely halving the chromosome complement. Crossing over between homologous chromosomes (homologs) is essential for their accurate segregation and defects are associated with infertility, miscarriage, and congenital disease. Factors that ensure crossing over between each pair of homologs include mammalian RING-domain proteins RNF212, HEI10, and RNF212B, alleles of which are linked to infertility and heritable variation in crossover rate. This study focuses on understanding the functions and relationships between pro-crossover RING proteins (CORs) in mouse, providing important insights into their roles in regulating recombination, the DNA repair process that gives rise to crossovers. Notably, chromosomal localization dynamics of the three CORs are distinct and show striking sexual dimorphism with important implications for models of crossover control.

## INTRODUCTION

During sexual reproduction, parents contribute equally to their offspring by producing gametes with precisely half the normal cellular ploidy. This is achieved via the two successive rounds of chromosome segregation that occur during meiosis. Accurate segregation during the first meiotic division (MI) requires crossing over between each pair of homologous chromosomes (homologs). In combination with cohesion between sister-chromatids, crossovers create connections called chiasmata that enable the stable biorientation of homolog pairs on the meiosis-I spindle ^1–3^. Defective crossing over can trigger cell death via the spindle checkpoint ^4^, or cause aneuploidy due to chromosome missegregation ^5, 6^, and is therefore associated with infertility, miscarriage, and congenital syndromes such as Down, Turner and Klinefelter ^7–10^.

Crossing over is typically the minority outcome of meiotic recombination with most events being repaired as noncrossovers without exchange of chromosome arms; for example, in mouse spermatocytes only ∼24 crossovers are selected from ∼250 recombination events initiated by programmed DNA double-strand breaks (DSBs) ^11^. Crossover sites are selected in such a way that each pair of chromosomes almost always obtains at least one crossover, termed crossover assurance; adjacent events tend to be widely and evenly spaced via crossover interference (which also limits total crossovers); and crossover numbers per nucleus show low variance relative to DSBs, a buffering process known as crossover homeostasis ^2, 3, 12^. The molecular basis of crossover patterning remains unclear but manifests at the cytological level as the selective retention/accumulation of certain pro-crossover factors at designated crossover sites as prophase progresses ^2, 3^. Among these are RING-domain proteins related to budding yeast Zip3 ^13^, inferred to mediate protein modification by ubiquitin and the related small-protein modifier SUMO ^2, 14–21^. Species differ with respect to the number and subgroup (Zip3/RNF212 and HEI10) of these **C**ross**O**ver **R**ING proteins (CORs). For example, *Sordariales* and plant genomes encode a single HEI10 homolog ^13, 18, 22^; *Drosophila* encodes three Zip3/RNF212-like proteins, two of which appear to be redundant ^23^ ^21^; and *C. elegans* has four members with homology to both subgroups ^19, 20, 24^.

Mammalian genomes encode both RNF212 and HEI10 (a.k.a. CCNBIP1) homologs with interrelated functions that are essential for crossing over ^15–17, 25^. RNF212 specifically localizes between synapsed chromosomes, initially as numerous small foci all along the central region of synaptonemal complexes (SCs), before undergoing HEI10-dependent repatterning to accumulate at prospective crossover sites while being lost elsewhere ^15–17^. RNF212 and other CORs are inferred to stabilize pro-crossover factors that directly mediate the DNA events of crossing over ^15–19, 26, 27^. These include the MutSγ complex, a heterodimer of MSH4 and MSH5 that can bind D-loop and Holliday junction intermediates ^28–30^, and is inferred to stabilize and protect them from dissociation by the conserved Bloom-helicase/decatenase complex, BLM-TOPIIIα-RMI1-RMI2 (analogous to Sgs1-Top3-Rmi1 in budding yeast)^31–34^. In mouse spermatocytes, MutSγ initially localizes to a majority of recombination sites during late zygotene and early pachytene as homologs complete synapsis ^11, 15^. Subsequently, MutSγ is lost from most sites but retained at prospective crossover sites, which then accumulate crossover-specific factors such as the MutLγ endonuclease ^2, 29^.

A key question for understanding meiotic crossover control is whether the dynamic focal patterning of COR proteins along SCs is responsible for specifying crossover sites, as proposed by recent models that invoke phase-transition/condensation, coarsening, and/or Ostwald ripening processes ^19, 35–37^. In these models, adjacent recombination sites compete to accumulate a limiting amount of COR protein that diffuses along synaptonemal complexes, with crossover designation ensuing at “winning” sites where COR has been accrued. Alternatively, COR patterning may occur as a downstream consequence of crossover patterning by a distinct process such as the mechanical stress envisioned in the beam-film model ^38^ ^39^. The latter possibility does not exclude COR coarsening, which could help reinforce initial crossover designation or have other downstream functions.

Here we characterize mammalian CORs RNF212, HEI10, and a new member RNF212B ^40–45^. Contrasting phenotypes, localization, and genetic dependencies imply that RNF212B, RNF212, and HEI10 function as distinct modules that integrate signals from DSBs, synapsis, the cell-cycle, and developing crossover complexes in order to effect the spatiotemporal coordination of the major events of meiotic prophase-I. Moreover, the three CORs show diverse spatiotemporal dynamics, including striking differences between males and females, features that are not readily reconciled by coarsening models of crossover patterning.

## RESULTS

### *Rnf212b^-/-^* mutant mice are sterile due to diminished crossing over

Null mutations in mouse *Rnf212b* were created by pronuclear injection with CRISPR-Cas9, targeting sequences immediately downstream of the first ATG (**Figures S1** and **S2**). Seven mutated lines were obtained, the mutations and phenotypes of which are summarized in **Table S1**. Two identical mutant lines, with a single nucleotide insertion that results in a frameshift after the eighth codon and a premature stop codon after just 27 codons (**Figure S2B**), were chosen for detailed phenotypic analysis. An independent *Rnf212b* mutant mouse line was recently reported with similar phenotypes to those described below ^45^.

Heterozygous mutant animals were backcrossed four times and then bred to homozygosity. Mature *Rnf212b^-/-^* mutant males had testes that were ∼3-fold smaller than wild-type counterparts (means ± SDs of 219.8 ± 16.8 mg per pair of testes from 8 wild-type mice versus 71.6 ± 7.8 mg from 6 *Rnf212b^-/-^* mice; *p* < 0.0001, two-tailed *t* test; **Figure 1A**), the cauda epididymides were devoid of sperm, and the animals were sterile. Testis sections from *Rnf212b^-/-^* mutant males revealed that post-metaphase-I stages and spermatozoa were absent indicating loss of spermatocytes at stage XII (**Figure 1B**). Sterility was also observed for female *Rnf212b^-/-^* mutants, analysis of which is presented in **Figure 6**.

**Figure 1.**
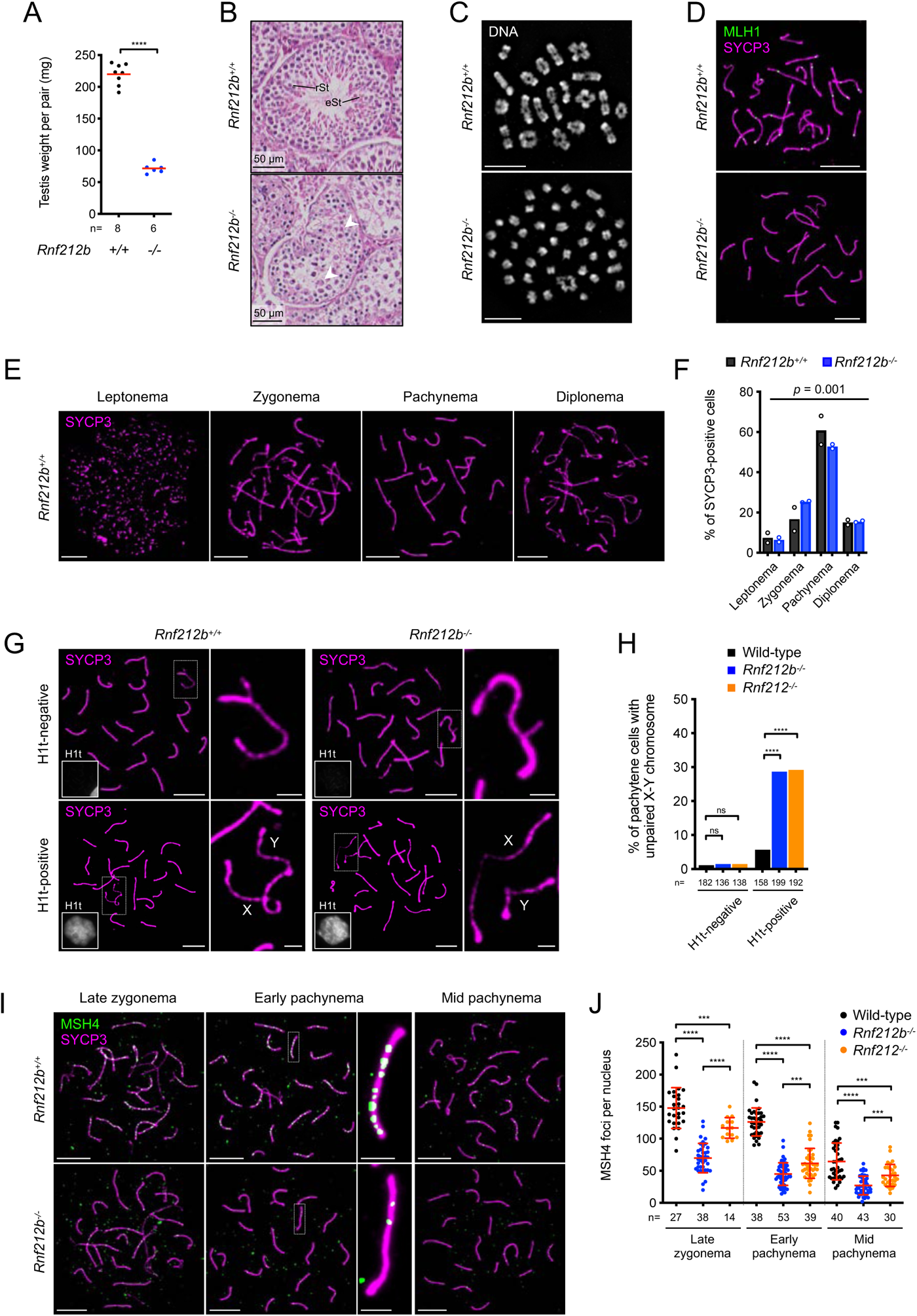
RNF212B is required for crossing over via stabilization of ZMMs in mouse spermatocytes. (A) Reduced testis size in *Rnf212b^-/-^*mutant males. Red bars indicate means. \*\*\*\**p* ≤ 0.0001 for a two-tailed *t* test. n, numbers of mice analyzed. (B) Defective spermatogenesis in *Rnf212b^-/-^* mutant males. Seminiferous tubule sections from wild-type and *Rnf212b^-/-^* testes stained with hematoxylin and eosin. rSt, round spermatid; eSt, elongated spermatid. Arrowheads indicate metaphase cells. (C) Chiasmata are diminished in *Rnf212b^-/-^* spermatocytes. Diakinesis/metaphase-I stage spermatocytes from wild-type and *Rnf212b^-/-^* testes stained with DAPI. (D) MLH1 foci are absent in *Rnf212b^-/-^* spermatocytes. Mid-late pachytene-stage spermatocyte nuclei from wild-type and *Rnf212b^-/-^* testes immunostained for SYCP3 and MLH1. (E) Representative images of successive prophase-I stages in chromosome spreads of wild-type spermatocytes immunostained for SYCP3. (F) Distributions of spermatocytes in successive prophase-I stages based on SYCP3 staining. >250 unselected SYCP3-positive cells were analyzed per mouse. Bar graphs indicate means of two mice with plots for each mouse. *p* = 0.001 for *G* test (total cells analyzed: 578 *Rnf212b^+/+^* cells and 549 *Rnf212b ^-/-^*cells). (G) Autosomal and X-Y synapsis in wild-type and *Rnf212b^-/-^* spermatocytes. Pachytene-stage spermatocyte nuclei immunostained for SYCP3 and H1t are shown. The magnified images show X-Y chromosomes. (H) Premature desynapsis of X-Y chromosomes in *Rnf212b^-/-^* and *Rnf212^-/-^* spermatocytes. ns, not significant (*p* > 0.05); \*\*\*\**p* ≤ 0.0001 for Fisher’s exact tests. Total numbers of cells analyzed from 3 mice of each genotype are indicated below the X axis. (I) Wild-type and *Rnf212b^-/-^* spermatocyte nuclei at the indicated stages immunostained for SYCP3 and a ZMM factor MSH4. Representative chromosomes are magnified. (J) MSH4 focus counts. Red bars indicate means ± SDs. \*\*\**p* ≤ 0.001; \*\*\*\**p* ≤ 0.0001 for two-tailed Mann-Whitney tests. n, numbers of nuclei analyzed. Scale bars, 10 μm for images of full nuclei and 2 μm for magnified panels.

*Rnf212^-/-^* mutants are sterile because crossing over almost completely fails, resulting in unconnected univalent chromosomes at metaphase I ^15^. To determine whether this is also the case for *Rnf212b^-/-^* mutants, chromosome spreads from diakinesis/metaphase-I spermatocytes were analyzed (**Figure 1C**). In wild-type nuclei, all autosomes presented as bivalents with 24.1 ± 2.3 chiasmata per nucleus (mean ± SD; *n =* 61 nuclei). Conversely, *Rnf212b*^-/-^ nuclei contained primarily univalents with a residual of 1-3 chiasmata (mean ± SD; 0.66 ± 0.75 chiasmata, *n* = 77 nuclei). Immunostaining of surface-spread pachytene-stage chromosomes revealed that *Rnf212b*^-/-^ mutant spermatocytes completely failed to form foci of the crossover-specific marker MLH1 (**Figure 1D**), indicating defects in the designation and/or maturation of crossover-specific recombination complexes. This inference was confirmed by analyzing two other crossover markers, PRR19 and CDK2 (**Figure S3**) ^16, 45, 46^.

### Irregular chromosome synapsis in *Rnf212b^-/-^* mutant spermatocytes

To begin to understand the defects leading to crossover failure in the *Rnf212b^-/-^* mutant, progression through meiotic prophase-I was assessed by quantifying the fractions of spermatocytes at each substage (leptonema, zygonema, pachynema and diplonema; **Figure 1E**). Stage distributions indicated a significant overrepresentation of zygotene-stage nuclei and fewer pachytene nuclei in *Rnf212b^-/-^* mutant spermatocytes relative to wild type (*p* = 0.001, *G* test) suggesting delayed or unstable synapsis (**Figure 1F**). Consistently, X-Y chromosomes were frequently unsynapsed in mutant spermatocytes, although they were always in close proximity (**Figure 1G** and **1H**). Unsynapsed X-Y chromosomes were over 5-fold more frequent in *Rnf212b^-/-^* spermatocytes relative to wild type (28.6%, *n* = 57/199 cells from 3 mice and 5.7%, *n* = 9/158 cells from 3 mice, respectively; *p* < 0.0001, Fisher’s exact test) and were observed specifically in mid/late pachytene cells (staining positive for the histone variant H1t) indicating premature X-Y de-synapsis analogous to that seen in the *Rnf212^-/-^* mutants ^15^.

### Intermediate steps of meiotic recombination are defective in *Rnf212b^-/-^* mutants

The modest synapsis defects of the *Rnf212b^-/-^* mutant suggest that early steps of recombination may occur normally, with RNF212B promoting later steps of crossover designation and/or maturation. To test these inferences, a variety of recombination markers were analyzed by immunostaining spermatocyte chromosome spreads (**Figures 1I**, **S4** and **S5**). RAD51 and DMC1 assemble onto resected DSB ends to form nucleoprotein filaments that catalyze homologous pairing and DNA strand exchange ^47^. The numbers of RAD51 and DMC1 foci detected in zygotene spermatocytes were slightly elevated in the *Rnf212b^-/-^* mutant relative to wild type (**Figure S4**), suggesting a mild perturbation of recombination as chromosomes synapse that might reflect the delayed/unstable synapsis inferred above (**Figure 1G,H**).

Replication protein A (RPA; comprising RPA1, RPA2 and RPA3) binds single-stranded DNA formed at DSB ends and at D-loops as strand exchange ensues ^48^. Prominent immunostaining foci of RPA2 emerge in zygonema, as DMC1 and RAD51 foci are diminishing, and then disappear as DSB repair ensues during pachynema. In mid-zygonema, *Rnf212b^-/-^* and wild-type spermatocytes formed similar numbers of RPA2 foci, but in subsequent stages levels were significantly lower in *Rnf212b^-/-^* cells suggesting that DSBs are repaired faster in the mutant (**Figure S5A,B**). More dramatic reductions were observed for foci of the ZMM proteins MSH4, TEX11^Zip4^ and HFM1^Mer3^, with ∼3-fold fewer foci in the *Rnf212b^-/-^* mutant (**Figures 1I,J** and **S5C-F**). The altered dynamics of RPA2 and ZMM factors in *Rnf212b^-/-^* mutant spermatocytes were similar to those seen for *Rnf212^-/-^*, although reduction of MSH4 foci appears to be more severe in *Rnf212b^-/-^* cells (**Figures 1I,J** and **S5**) ^15^.

### RNF212B dynamically localizes to synaptonemal complexes and recombination sites

To address the possibility that RNF212B functions locally to stabilize ZMMs at nascent crossover sites, antibodies were raised against full-length RNF212B (isoform a; **Figure S1**) and used to immunostain surface-spread spermatocyte chromosomes (**Figures 2A**,**B** and **S6**). RNF212B was first detected in very early zygonema, specifically localizing to sites of synapsis initiation marked by SYCP1 (73.1% of short SYCP1 stretches <2 μm colocalized with RNF212B foci; *n =* 38/52 from 11 nuclei). As synapsis ensued, a punctate staining pattern emerged along extending SCs, with focus numbers peaking in early pachynema, just after synapsis was completed, at 175.9 ± 15.2 foci per nucleus (mean ± SD; *n =* 27 nuclei; **Figure 2B**). Focus numbers then diminished throughout pachynema, but at the same time a small number of large, bright “amplified” RNF212B foci emerged. By late pachynema, each SC had only one or two large RNF212B foci with 25.0 ± 2.3 foci per nucleus (mean ± SD; *n =* 29 nuclei; **Figure 2B**). By mid-diplonema, as homologs desynapsed, all RNF212B foci had disappeared.

**Figure 2.**
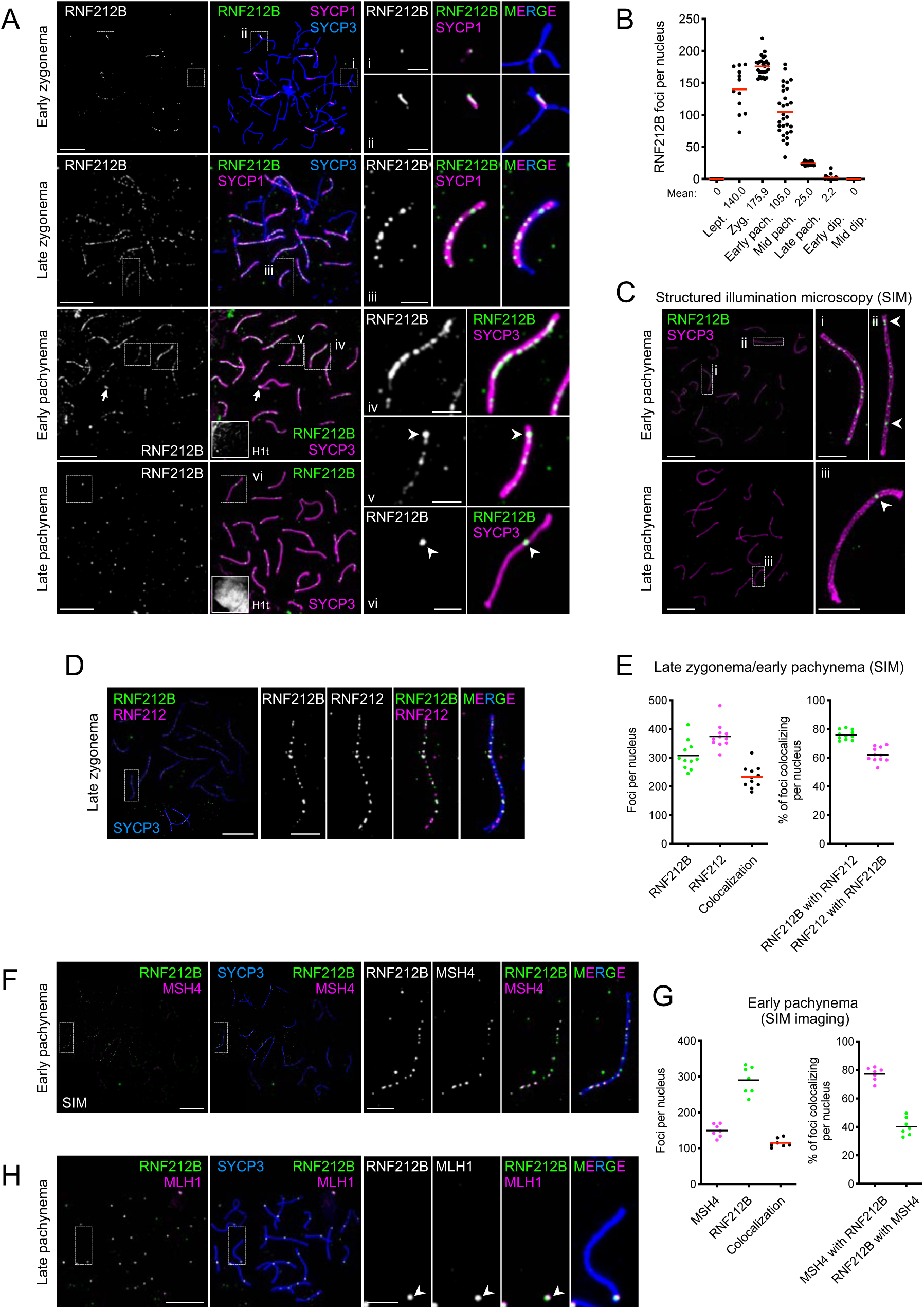
RNF212B localization to synaptonemal complexes and recombination sites in prophase I spermatocytes. (A) RNF212B localization in spermatocyte nuclei at successive prophase I stages. Images show surface-spread prophase-I spermatocyte nuclei immunostained for SYCP3, SYCP1, RNF212B and H1t. Arrowheads highlight large RNF212B foci. The arrow indicate synapsed pseudo-autosomal region between the X-Y chromosomes. (B) Numbers of RNF212B foci per nucleus in spermatocytes at successive prophase I stages. Red bars indicate means. Means of focus numbers are shown below the graph. Lept., leptonema; Zyg., zygonema; pach., pachynema; dip., diplonema. Numbers of nuclei analyzed were 8, 12, 27, 29, 29, 19 and 11 for in leptonema, zygonema, early pachynema, mid pachynema, late pachynema, early diplonema and mid diplonema, respectively. (C) RNF212B localizes to the central region of the synaptonemal complex. SIM images of early and late pachytene spermatocytes immunostained for SYCP3 and RNF212B. Arrowheads highlight large RNF212B foci. (D) RNF212B colocalization with RNF212 in late zygonema. SIM images of a late zygotene spermatocyte immunostained for SYCP3, RNF212B and RNF212. (E) Quantification of RNF212B colocalization with RNF212. Left, focus counts. Right, degree of colocalization. Black and red bars indicate means. 3 late-zygotene and 8 early-pachytene nuclei were analyzed by SIM imaging. (F) RNF212B colocalization with MSH4 in early pachynema. SIM images of an early pachytene spermatocyte immunostained for SYCP3, RNF212B and MSH4. (G) Quantification of RNF212B colocalization with MSH4. Left, focus counts. Right, degree of colocalization. Black and red bars indicate means. 7 early pachytene nuclei were analyzed by SIM imaging. (H) RNF212B colocalization with MLH1. A late pachytene spermatocyte immunostained for SYCP3, RNF212B and MLH1. Arrowheads indicate crossover sites. Magnified images in (A), (C), (D), (F), and (H) show representative chromosomes. Scale bars,10 μm for full nuclei and 2 μm for magnified images.

Super-resolution structured illumination microscopy (SIM) showed that RNF212B, like RNF212, localizes specifically to the central region of the SC (**Figure 2C**). The phenotypes of *Rnf212b^-/-^* mutants and the localization dynamics of the RNF212B protein indicate strong similarities with RNF212. To begin to explore the relationship between RNF212B and RNF212, colocalization was analyzed using SIM (**Figure 2D,E**). In late zygotene/early pachytene stage spermatocytes, when focus numbers peaked, an average of 307.5 ± 50.0 RNF212B foci were detected per nucleus compared to 374.6 ± 43.2 RNF212 foci, and 233.4 ± 38.7 RNF212B-RNF212 co-foci. Thus, RNF212-RNF212B colocalization is high but incomplete with 75.9 ± 3.4% of RNF212B foci colocalized with RNF212, whereas 62.1 ± 5.1% of RNF212 foci colocalized with RNF212B (means ± SDs; *n =* 11 nuclei; **Figure 2D,E**; note that many more foci are resolved by SIM than by the conventional fluorescence microscopy used in **Figure 2A,B**).

SIM was also used to analyze localization of RNF212B to ongoing recombination sites marked by ZMM factor MSH4 (**Figure 2F,G**). In early pachynema, RNF212B were in 2-fold excess relative to MSH4 (290.0 ± 38.3 RNF212B versus 149.4 ± 17.8 MSH4 foci per nucleus; means ± SDs; *n =* 7 nuclei). Reflecting this excess, only 40.2 ± 6.2 % of RNF212B foci colocalized with recombination sites marked by MSH4 foci; however, 77.2 ± 4.7% MSH4 foci colocalized with RNF212B (means ± SDs; *n =* 7 nuclei; **Figure 2F,G**).

In late pachynema, numbers of amplified RNF212B foci per nucleus were indistinguishable from those of the crossover marker MLH1 and colocalization was essentially absolute (**Figure 2H**; 99.7 ± 1.0 % of RNF212B foci colocalized with MLH1 foci and 99.0 ± 1.7 % of MLH1 foci colocalized with RNF212B foci; means ± SDs; *n =* 15 nuclei). Thus, RNF212B foci in late-pachytene spermatocytes exclusively localize to crossover sites.

### Amplified RNF212B foci and HEI10 designate crossover sites by mid pachynema

We previously showed that immunostaining foci of the ubiquitin ligase HEI10 localize to designated crossover sites marked by MLH1 foci in late pachynema ^16, 17^. Unlike RNF212B and RNF212, we did not detect numerous HEI10 foci in zygotene and early pachytene-stage spermatocytes. Instead, crossover specific HEI10 foci materialized in pachytene, similar to other crossover markers such as MLH1 (**Figure 3A,B**). By sub-staging pachytene spermatocytes, we showed that initial emergence of MLH1 foci occurred only in nuclei with a full complement of HEI10 foci, which was already present by mid pachynema (**Figure 3A,B**), i.e. crossover designation must have occurred by this stage.

**Figure 3.**
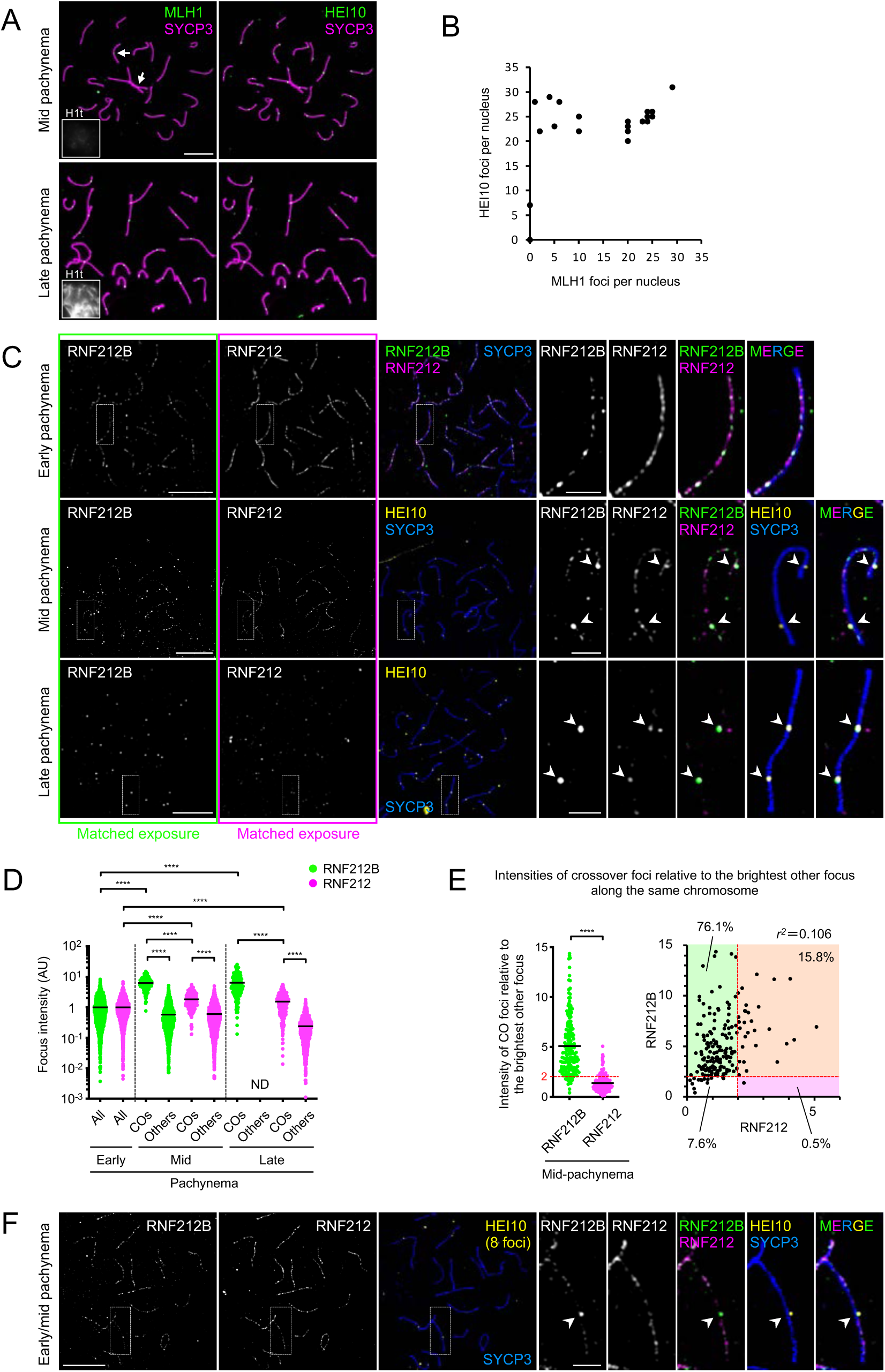
Distinct timing and development of crossover-specific foci of HEI10, RNF212B and RNF212. (A) HEI10 forms crossover foci by mid-pachynema. Pachytene spermatocytes nucleiimmunostained for SYCP3, HEI10, MLH1 and H1t. Top images show a representative H1t-negative, mid-pachytene nucleus in which small MLH1 foci are emerging on just a few chromosomes (indicated by arrows) whereas bright HEI10 foci are present on all autosomes. Bottom images show a representative H1t-positive late-pachytene nucleus in which both MLH1 and HEI10 form clear, overlapping crossover-specific foci on all autosomes. (B) Quantification of HEI10 and MLH1 foci in pachytene spermatocytes confirms that MLH1 foci develop only in nuclei with a full complement of HEI10 foci. 25 pachytene nuclei were randomly selected and analyzed. (C) Differentiation of RNF212B at prospective crossover sites is stronger than RNF212. Airyscan images of early, mid and late pachytene spermatocytes nuclei immunostained for SYCP3, RNF212B, RNF212 and HEI10. In all cases, RNF212B and RNF212 image exposures are matched. Arrowheads indicate crossover sites marked by HEI10. (D) Quantification of focus intensities for RNF212B and RNF212. Black bars indicate means. \*\*\*\**p* ≤ 0.0001 for two-tailed Mann-Whitney tests. 8 nuclei for each stage were analyzed by Airyscan imaging. Total numbers of foci analyzed: 2,449 RNF212B foci and 2,513 RNF212 foci in early pachynema; 194 crossover foci, 2,004 other RNF212B foci and 2,291 other RNF212 foci in mid pachynema; 206 crossover foci and 988 other RNF212 foci in late pachynema. All, all foci; COs, crossover foci colocalized with HEI10 foci; Others, other foci that don’t colocalize with HEI10 foci. ND, not detected. Quantification of focus intensities in representative individual nuclei in (D) is shown in **Figure S7A**. (E) Per chromosome analysis for autosomes in mid-pachytene spermatocyte nuclei. Intensity of each crossover-associated RNF212B or RNF212 focus relative to that of the brightest other (non-crossover) focus along the same chromosome plotted separately (left) and in two dimensions (right). Red dashed lines indicate a relative focus intensity of 2. Detailed representation of a per chromosome analysis is shown in **Figure S7B**. \*\*\*\**p* ≤ 0.0001 for two-tailed Mann-Whitney tests. 186 crossover foci from 8 mid-pachytene nuclei were analyzed. Two crossover-associated RNF212B foci with no other RNF212B foci along the same chromosome were excluded from the plots. (F) RNF212B differentiation without detectable RNF212 differentiation during early to mid pachyenema. Airyscan images of an early- to mid-pachytene spermatocyte nucleus, with 8 HEI10 foci, immunostained for SYCP3, RNF212B, RNF212 and HEI10 are shown. Arrowheads indicate crossover sites marked by HEI10. Magnified images in (C) and (F) show representative chromosomes. Scale bars, 10 μm for full nuclei and 2 μm for magnified images.

Having defined HEI10 foci as the earliest detectable marker of crossover designation, we set out to determine the spatial-temporal relationship with amplification of RNF212B and RNF212 foci. The intensity of each RNF212B or RNF212 focus was measured from images with matched exposures and normalized to the mean intensities of foci in early pachytene nuclei in which HEI10 foci were not yet detected (*n =* 2,449 RNF212B foci and 2,513 RNF212 foci; 8 nuclei; **Figure 3C**,**D** and **S7A**). In mid-pachytene nuclei with at least one clear HEI10 crossover focus on each autosomal SC, RNF212B foci at crossover sites (i.e. colocalizing with HEI10 foci) were 11-fold brighter than other RNF212B foci in the same nuclei (i.e. the average of all other foci from the same 8 nuclei; 6.29 ± 2.66 for 194 crossover foci versus 0.57 ± 0.59 for 2,004 other foci from 8 nuclei; means ± SDs; *p* < 0.0001, Mann-Whitney test) and 6.3-fold brighter than the foci detected in early pachytene nuclei, consistent with the interpretation that crossover foci grow at the expense other foci. The same analysis showed that RNF212 foci located at crossover sites were only 3-fold brighter than other foci in the same nuclei (1.83 ± 1.05 for 194 crossover foci versus 0.60 ± 0.58 for 2,291 other foci form 8 nuclei; means ± SDs; *p* < 0.0001, Mann-Whitney test) and 1.8-fold brighter than those in early pachytene. Thus, at designated crossover sites in mid pachytene, RNF212B foci show stronger amplification than RNF212 foci.

In late pachytene spermatocytes, the only remaining RNF212B foci were crossover specific and had intensities comparable to those of the crossover foci analyzed in mid-pachytene nuclei (6.42 ± 4.84-fold brighter than early pachytene foci; 206 crossover foci from 8 nuclei; mean ± SD) implying that RNF212B foci do not continue to enlarge between mid and late pachytene; and that RNF212B is lost from other sites as opposed to being absorbed into crossover foci. By contrast, RNF212 staining in late pachytene nuclei comprised a mixture of crossover foci and other small residual foci. The crossover-associated RNF212 foci in these nuclei were 6-fold brighter than other foci in the same nuclei (1.53 ± 1.13 for 206 crossover foci versus 0.24 ± 0.23 for 988 other foci from 8 nuclei; means ± SDs; *p* < 0.0001, Mann-Whitney test) but only 1.5-fold brighter than those in early pachytene. Thus, crossover-associated RNF212 foci continue to differentiate between mid and late pachynema but this may occur primarily via loss of RNF212 from noncrossover sites as opposed to continued growth of crossover-specific foci. Consistent with these inferences, the total signal intensity of RNF212B per nucleus was unchanged from early to mid pachynema, but then decreased from mid to late pachynema as RNF212B was lost from noncrossover sites; by comparison, total signal intensity of RNF212 reduced during both early to mid, and mid to late pachynema (1.00 ± 0.21, 0.97 ± 0.24, and 0.54 ± 0.33 for RNF212B; and 1.00 ± 0.32, 0.69 ± 0.31, and 0.22 ± 0.10 for RNF212 in early, mid, and late pachynema, respectively, *n =* 8 nuclei each; **Figure S7C**).

Per chromosome analysis for mid-pachytene autosomes highlighted additional distinctions between RNF212B and RNF212 with respect to the amplification of brighter crossover-correlated foci (**Figures 3E** and **S7B**). RNF212B foci at crossover sites were on average 5.1-fold brighter relative to the brightest other focus along the same chromosome, whereas RNF212 foci were only 1.4-fold brighter (**Figure 3E**, left). When these relative intensities were plotted for individual pairs of colocalizing RNF212-RNF212B crossover foci, almost no correlation was detected (*r*^2^ = 0.106), i.e. at mid pachynema, bright RNF212B foci are not coincident with bright RNF212 foci, pointing to distinct amplification behaviors of the two proteins at designated crossover sites (**Figure 3E**, right).

In mid-pachynema, 76.1% of designated crossover sites (*n =* 140/184; 8 nuclei) showed amplification for RNF212B foci (≥2-fold brighter relative to the brightest other focus along the same chromosome) but not for RNF212 foci. Oppositely only 0.5% of crossover sites (*n =* 1/184; 8 nuclei) showed amplification of only RNF212 but not RNF212B (**Figure 3E**, right) Together, these data imply that crossover-specific amplification of RNF212B occurs earlier than that of RNF212. The distinct behaviors of RNF212B and RNF212 were most pronounced in nuclei transitioning between early and mid-pachynema, as HEI10 foci were emerging (i.e. nuclei with <19 HEI10 foci per nucleus; **Figure 3F**). In these nuclei, RNF212B showed clear amplification at designated crossover sites but RNF212 did not.

Together, our analysis of focus dynamics in spermatocytes shows that RNF212B foci amplify at designated crossover sites during early-to-mid pachytene, coincident with the appearance of HEI10 foci (**Figure S8**); and then RNF212B is lost from other sites without additional amplification of crossover-associated foci. By contrast, RNF212 crossover foci differentiate later, during mid-to-late pachytene; and loss of RNF212 from other sites, rather than growth, may be the major cause of changes in relative intensity.

### Interdependence between HEI10, RNF212B, and RNF212

Our previous studies evoked a model in which RNF212 establishes a precondition for crossover/noncrossover differentiation by stabilizing nascent recombination intermediates and rendering further progression dependent on HEI10 ^16, 17^. Notably, in *Hei10^mei4/mei4^* mutant spermatocytes, abundant RNF212 foci persist along synapsed chromosomes and recombination stalls at an intermediate step marked by the persistence of ZMM foci throughout pachytene ^16, 17^. Similarly, in early pachynema *Hei10^mei4/mei4^* mutant spermatocytes, initial RNF212B staining was indistinguishable from wild type, but this pattern of abundant foci along SCs persisted until early diplonema (**Figure 4A,B**). Moreover, in sharp contrast to the crossover-specific pattern of amplified RNF212B foci that developed in wild-type nuclei (24.8 ± 2.7 foci, mean ± SD from 16 nuclei), significantly amplified RNF212B foci were not detected along the majority of chromosomes in *Hei10^mei4/mei4^* mutant nuclei (**Figure 4C,D**). Specifically, for 54.4% of individual chromosomes from late pachytene *Hei10^mei4/mei4^* spermatocyte nuclei (marked by strong H1t staining), neither the first nor the second brightest focus was ≥1.5-fold brighter than the third brightest focus along the same chromosome, and only 17.0% of individualchromosomes had RNF212B foci that were ≥2-fold brighter than the third brightest focus along the same chromosome (*n =* 93/171 and 29/171 chromosomes, respectively; 9 nuclei; **Figure 4A,C**). The most differentiated RNF212B foci were on average only 1.6-fold brighter (with a maximum of 4.2-fold) relative to the third brightest focus along the same chromosome in late pachytene *Hei10^mei4/mei4^* spermatocyte nuclei; whereas in wild-type mid-pachytene nuclei, RNF212B foci at crossover sites were on average 6.7-fold brighter relative to the brightest other focus along the same chromosome (**Figure 4D**; 33 wild-type chromosomes with two crossover foci were used for comparison to *Hei10^mei4/mei4^* chromosomes; 8 nuclei). Thus, crossover-specific patterning of RNF212B appears to be largely dependent on HEI10. Reciprocally, immunostaining for HEI10 in *Rnf212b^-/-^* mutant spermatocytes revealed that formation of HEI10 foci is dependent on RNF212B (**Figure 4E**).

**Figure 4.**
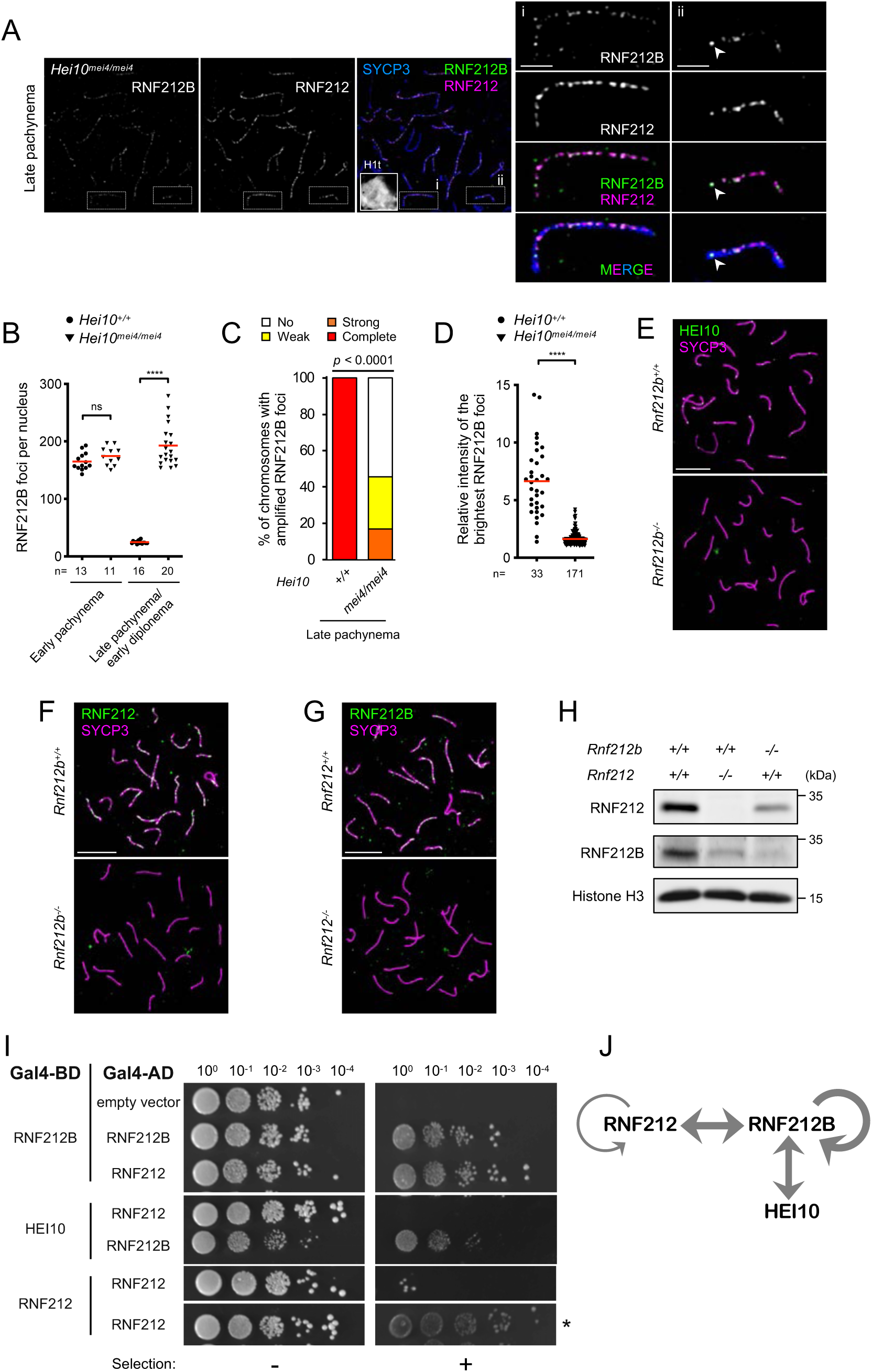
Interdependent relationship between HEI10, RNF212 and RNF212B. (A) Persistent RNF212B staining in *Hei10^mei4/mei4^* spermatocytes. Images of a late-pachytene nucleus immunostained for SYCP3, RNF212B, RNF212 and H1t. The magnified panels show representative chromosomes with: (i) no apparent differentiation of RNF212B foci; or some differentiation, indicated by arrowheads. Scale bars, 10 μm in images of full nuclei and 2 μm in the magnified panels. (B) Focus counts of RNF212B. Red bars indicate means. Total numbers of nuclei analyzed are indicated below the X-axis. For late stages, only late pachytene nuclei were analyzed for *Hei10^+/+^* whereas both late pachytene and early diplotene nuclei were analyzed for *Hei10^mei4/mei4^* because of the early desynapsis of *Hei10^mei4/mei4^* cells ^16^. ns, not significant (*p* > 0.05); \*\*\*\**p* ≤ 0.0001 for two-tailed Mann-Whitney tests. (C) Differentiation of RNF212B foci. Autosomes in late-pachytene *Hei10^+/+^* and *Hei10^mei4/mei4^* spermatocyte nuclei were categorized into four classes based on the degree of RNF212B differentiation. Strong, weak or no differentiation represents autosomes where the brightest RNF212B foci were ≥2-fold, 1.5-2-fold, or <1.5-fold brighter relative to the third brightest focus along the same synaptonemal complex, respectively. Complete differentiation is autosomes that have only one or two bright RNF212B foci. 8 late-pachytene *Hei10^+/+^* and 9 late-pachytene *Hei10^mei4/mei4^* nuclei were analyzed by Airyscan imaging. *p* < 0.0001 for a *G* test. (D) Intensity of the brightest RNF212B foci relative to the third brightest focus along the same chromosome in late pachytene *Hei10^mei4/mei4^* and mid pachytene *Hei10^+/+^* spermatocyte nuclei. Red bars indicate means. Nuclei analyzed were the same as in (C) and Figure 3E, respectively, but chromosomes with two crossover foci were analyzed for *Hei10^+/+^* spermatocytes. Total numbers of foci analyzed are indicated below the X-axis. \*\*\*\**p* ≤ 0.0001 for two-tailed Mann-Whitney test. (E) Crossover-specific immunostaining foci of HEI10 foci were absent from chromosome spreads of *Rnf212b^-/-^* spermatocytes. Mid/late pachytene nuclei from wild-type and *Rnf212b^-/-^* spermatocytes were immunostained for SYCP3 and HEI10. Scale bars, 10 μm. (F and G) RNF212 (F) and RNF212B (G) are interdependent for chromosomal localization. Early pachytene spermatocyte nuclei from the indicated genotypes were immunostained for SYCP3 and RNF212 (F) or RNF212B (G). Scale bars, 10 μm. (H) Interdependent protein stability of RNF212B and RNF212. Whole-testis extract from indicated genotypes were subjected to immunoblotting for RNF212 and RNF212B. Histone H3 is a loading control. (I) Yeast two-hybrid assay showing interaction between RNF212B, RNF212 and HEI10, and self-interactions for RNF212B and RNF212. Yeast cells expressing Gal4 activation (Gal4-AD) and binding (Gal4-BD) domain fused to the indicated proteins, or empty vectors as controls, were spotted with the indicated dilutions. Aureobasidin A selection was used to detect activation of the *AUR1-C* reporter. * indicates less stringent selection for *ADE2* and *HIS3* reporters. (J) A summary of interactions between RNF212B, RNF212 and HEI10 detected by yeast two-hybrid assays in (I).

Our localization analysis is incompatible with RNF212B and RNF212 functioning as an obligate heterocomplex of fixed stoichiometry. Even so, immunostaining analysis in *Rnf212b^-/-^* and *Rnf212^-/-^* mutants indicated an interdependent relationship. RNF212 immunostaining was diminished in *Rnf212b^-/-^* mutant meiocytes and, likewise, RNF212B immunostaining was diminished in *Rnf212^-/-^* cells (**Figures 4F,G** and **S9**). Immunoblotting revealed that protein levels of RNF212 and, to a greater extent, RNF212B were reduced in testis extracts from *Rnf212b^-/-^* and *Rnf212^-/-^* mutants, respectively, relative to wild-type controls (**Figure 4H**).

Interaction between RNF212B and RNF212 was detected by yeast two-hybrid assay (Y2H), and self-interaction for both proteins was also inferred, although RNF212 self-interaction appeared to be much weaker than that of RNF212B (**Figure 4I,J**). Thus, RNF212B and RNF212 physically interact and are mutually dependent for protein stability and thus chromosomal localization. Additional analysis showed that an intact RING finger domain was essential for RNF212B self-interaction by Y2H, for the stability of RNF212B and RNF212 *in vivo*, and thus for the pro-crossover functions of RNF212B (**Figures S10** and **S11**). Interaction between HEI10 and RNF212B (but not HEI10 and RNF212) was also detected, which might reflect their spatiotemporal colocalization at designated crossover sites (**Figure 4I,J**).

### Similar but distinct localization of RNF212B and RNF212 in mutants defective for recombination and synapsis

The relationship between RNF212B and RNF212 was further explored by analyzing their localization in mutants defective for recombination or synapsis. Repair of SPO11-induced DSBs is required for the alignment and synapsis of homologs. Thus, *Spo11^-/-^* mutant meiocytes are generally defective for synapsis; however, extensive stretches of SC can form between non-homologous axes ^49, 50^. In *Spo11^-/-^* spermatocytes, RNF212B staining was still detected specifically on synapsed regions marked by SYCP1, as previously seen for RNF212 (**Figure 5A**)^15^. However, intensities of RNF212B immunofluorescence measured in *Spo11^-/-^* nuclei with late zygotene/early pachytene-like morphologies (well-developed SYCP3-staining axes and extensive synapsis, albeit non-homologous) were ∼50% lower than those in equivalent early-pachytene *Spo11^+/+^* nuclei (**Figure 5B**). By contrast, intensities of SC-associated RNF212 immunostaining were comparable for *Spo11^-/-^* and *Spo11^+/+^* nuclei. Correspondingly, the normalized ratio of RNF212B / RNF212 immunofluorescence intensity on synapsed regions was reduced by ∼50% in *Spo11^-/-^* spermatocytes (**Figure 5C**). Thus, SPO11-initiated recombination differentially influences the association of RNF212B and RNF212 with synaptonemal complexes, with only RNF212B showing a significant dependence.

**Figure 5.**
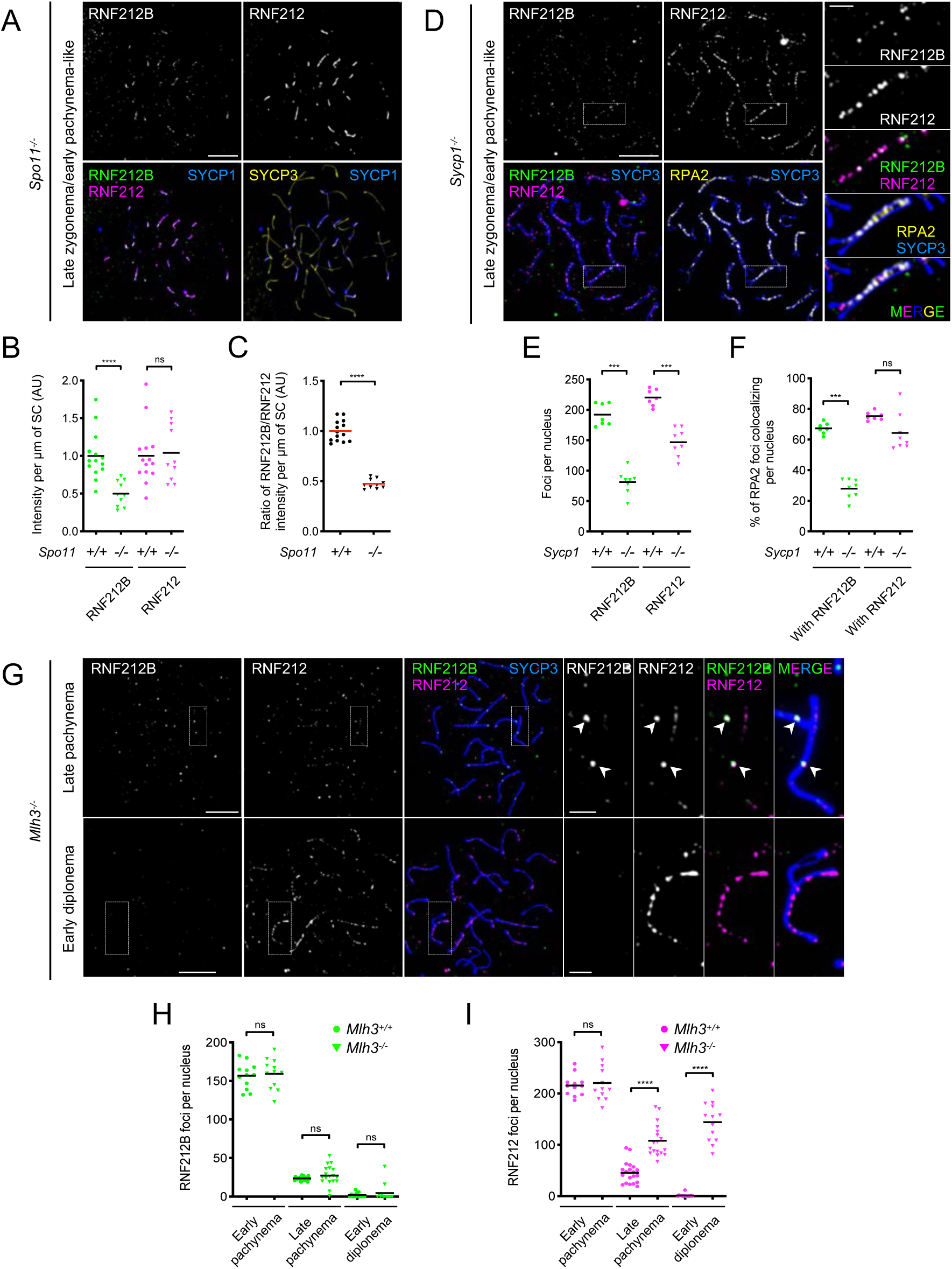
Chromosomal localization of RNF212B and RNF212 in mutants defective for recombination and synapsis. (A) RNF212B and RNF212 localize to synapsed regions between non-homologous axes in DSB-defective *Spo11^-/-^* spermatocytes. Images show a late zygotene/early pachytene-like nucleus immunostained for SYCP3, SYCP1, RNF212B and RNF212. (B and C) Staining intensities of RNF212B and RNF212 on synapsed regions. Signal intensities per μm of SC per nucleus (B); and ratios of intensities per μm of SC for RNF212B relative to RNF212 (C). Black and red bars indicate means. 14 early pachytene *Spo11^+/+^* nuclei and 10 late zygotene/early pachytene-like *Spo11^-/-^* nuclei were analyzed. (D) RNF212B and RNF212 localize to recombination sites between aligned homolog axes in synapsis-defective *Sycp1^-/-^* spermatocytes. A late zygotene/early pachytene-like nucleus immunostained for SYCP3, RNF212B, RNF212 and RPA2 is shown. (E and F) Focus counts of RNF212B and RNF212 (E); and degree of RPA2 colocalization with RNF212B and RNF212 (F). Black bars indicate means. 7 early pachytene *Sycp1^+/+^* nuclei and 8 late zygotene/early pachytene-like *Sycp1^-/-^* nuclei were analyzed. (G) Differentiation of crossover-specific foci of RNF212B and RNF212 and persistence of RNF212 staining in crossover-defective *Mlh3^-/-^* spermatocyte nuclei. Late pachytene and early diplotene nuclei immunostained for SYCP3, RNF212B, and RNF212 are shown. Arrowheads indicate large RNF212B foci. (H and I) Numbers of RNF212B (H) and RNF212 (I) foci per nucleus in successive late prophase I stages of spermatocytes. Black bars indicate means. Numbers of nuclei analyzed in early pachynema, late pachynema, and early diplonema were 12, 20, and 13 for *Mlh3^+/+^*; and 12, 18, and 13 for *Mlh3^-/-^*. ns, not significant (*p* > 0.05); \*\*\**p* ≤ 0.001; \*\*\*\**p* ≤ 0.0001 for two-tailed Mann-Whitney tests. Magnified images in (D) and (G) show representative chromosomes. Scale bars, 10 μm for full nuclei and 2 μm for magnified images.

SYCP1 is the major component of the SC central region. Although *Sycp1^-/-^* mutant meiocytes fail to assemble SCs, recombination initiates normally and homolog axes become coaligned ^51^. In *Sycp1^-/-^* spermatocytes, RNF212B staining was barely detectable in early/mid zygotene-like nuclei indicating that initial association with meiotic chromosomes is strongly dependent on the SC central region (**Figure S12A**). However, relatively dim RNF212B foci could be detected between aligned homologous axes in late zygotene/early pachytene-like nuclei that had extensive homolog coalignment (**Figures 5D**). The majority of these RNF212B foci localized to recombination sites, revealed by a high level of colocalization with RPA2 (71.0 ± 8.0%; mean ± S.D., *n =* 8 nuclei), as seen previously for RNF212 localization in *Sycp1^-/-^* spermatocytes (**Figures 5D** and **S12B**)^15^. Thus, normal localization of RNF212B to prophase-I chromosomes is strongly dependent on synapsis, with respect to both timing and abundance, but RNF212B can independently localize to recombination sites, albeit inefficiently.

RNF212B focus numbers in *Sycp1^-/-^* nuclei were reduced 2.4-fold relative to *Sycp1^+/+^* controls (192.1 ± 18.3 foci in *Sycp1^+/+^* versus 81.4 ± 19.3 in *Sycp1^-/-^*; mean ± SDs; **Figure 5E**). Correspondingly, the degree of RPA2 colocalization with RNF212B was reduced 2.4-fold (from 67.3 ± 3.7% in *Sycp1^+/+^* to 27.9 ± 6.0% in *Sycp1^-/-^*; means ± S.D.; *n* = 7 and 8 nuclei, respectively; *p* = 0.0002, Mann-Whitney test) and RNF212B-RPA2 co-foci were reduced 1.9-fold (109.4 ± 7.4 foci in *Sycp1^+/+^* versus 57.9 ± 14.7 in *Sycp1^-/-^*; mean ± SDs; **Figures 5F** and **S12C**). In contrast, in the same nuclei, numbers of RNF212-RPA2 co-foci remained essentially unchanged (122.4 ± 7.8 foci in *Sycp1^+/+^* versus 131.3 ± 17.4 in *Sycp1^-/-^*; mean ± SDs) despite a 1.5-fold reduction in total RNF212 focus numbers (220.3 ± 13.9 foci in *Sycp1^+/+^* versus 146.9 ± 21.4 in *Sycp1^-/-^*; mean ± SDs; **Figures 5E** and **S12C**). Moreover, the degree of RPA2-RNF212 colocalization was barely altered (75.2 ± 2.9% in *Sycp1^+/+^* versus 64.3 ± 11.6% in *Sycp1^-/-^*; mean ± SDs; *p* = 0.0578, Mann-Whitney test; **Figure 5F**).

We infer that the localization of RNF212B to recombination sites shows a greater dependence on synapsis than does RNF212. Consequently, this differential dependency resulted in reduced colocalization of RNF212B and RNF212 foci in *Sycp1^-/-^* spermatocytes (61.8 ± 9.0% of RNF212B foci colocalized with RNF212, and 33.9 ± 6.3% of RNF212 foci colocalized with RNF212B; versus 95.8 ± 2.4% and 83.8 ± 8.6% in *Sycp1^+/+^* nuclei; mean ± SDs; **Figure S12D**).

MLH3 and MLH1 constitute the MutLγ complex that facilitates the maturation and crossover-specific resolution of double-Holliday junctions ^29, 30^. Consistently, *Mlh3^-/-^* meiocytes are proficient for synapsis and early steps of meiotic recombination, but defective for crossing over^52^. We previously showed that MutLγ constrains HEI10 localization both temporally and spatially; in *Mlh3* mutant spermatocytes HEI10 foci appear much earlier than in wild type (during zygotene), are much more numerous (∼90 per nucleus), and persist throughout pachytene ^16^. In contrast, RNF212B staining in *Mlh3^-/-^* spermatocytes was comparable to *Mlh3^+/+^* controls: focus numbers peaked in early pachynema, then diminished as pachynema progressed with one or two large RNF212B foci emerging on each SC (**Figures 5G**,**H** and **S12E,F**). Numbers of amplified crossover-specific RNF212B foci in late pachytene *Mlh3^-/-^* nuclei were slightly lower than in wild-type *Mlh3^+/+^* controls, possibly reflecting reduced stability or altered efficiency of formation (**Figure S12F**). In addition, unlike wild type, some smaller foci remained on the SCs until late pachynema in *Mlh3^-/-^* nuclei suggesting an additional signal following the maturation of crossover sites may trigger the complete loss of RNF212B from noncrossover sites.

Nonetheless, this analysis indicates that the differentiation of RNF212B into crossover-specific foci precedes crossing over and does not require crossover-specific factor, MutLγ. Importantly, we can also infer that while crossover-specific patterning of RNF212B requires HEI10 (above, **Figure 4A-D**), it does not require HEI10 *per se* to undergo crossover-specific patterning, i.e. crossover-specific patterning of HEI10 appears to be downstream of initial crossover designation and requires a maturation step that is dependent on MutLγ.

RNF212 staining in *Mlh3^-/-^* mutant spermatocytes was comparable to wild-type cells in early pachynema, but the differentiation of crossover-specific foci was not as conspicuous; and overall focus numbers remained relatively high throughout pachynema (**Figures 5G**,**I**, and **S12G**). Moreover, numerous RNF212 foci persisted on synapsed regions in early diplotene *Mlh3^-/-^* nuclei in which RNF212B staining was barely detectable. Together, localization studies in mutant contexts extend our inference that the dynamics and regulation of RNF212B, RNF212, and HEI10 are distinct.

### RNF212B is essential for crossing over in oocytes

*Rnf212b^-/-^* mutation in females also caused sterility but unlike males, in which gametes were absent (**Figure 1B**), the ovaries of 18 days postpartum (dpp) mice contained large numbers of oocytes (**Figure 6A,B**). However, analysis of MLH1 foci in prophase-I nuclei (from fetal ovaries, **Figure 6C**), and chiasmata in metaphase-I oocytes (from adult ovaries, **Figure 6D**) revealed that a severe crossover defect is common to *Rnf212b^-/-^* mutants of both sexes: MLH1 foci were completely absent, and chiasmata were reduced over 20-fold (1.11 ± 0.92 chiasmata per nucleus in *Rnf212b*^-/-^, *n =* 38 oocytes; compared to 24.6 ± 2.4 in *Rnf212b*^+/+^, *n* = 28 oocytes; means ± SDs).

**Figure 6.**
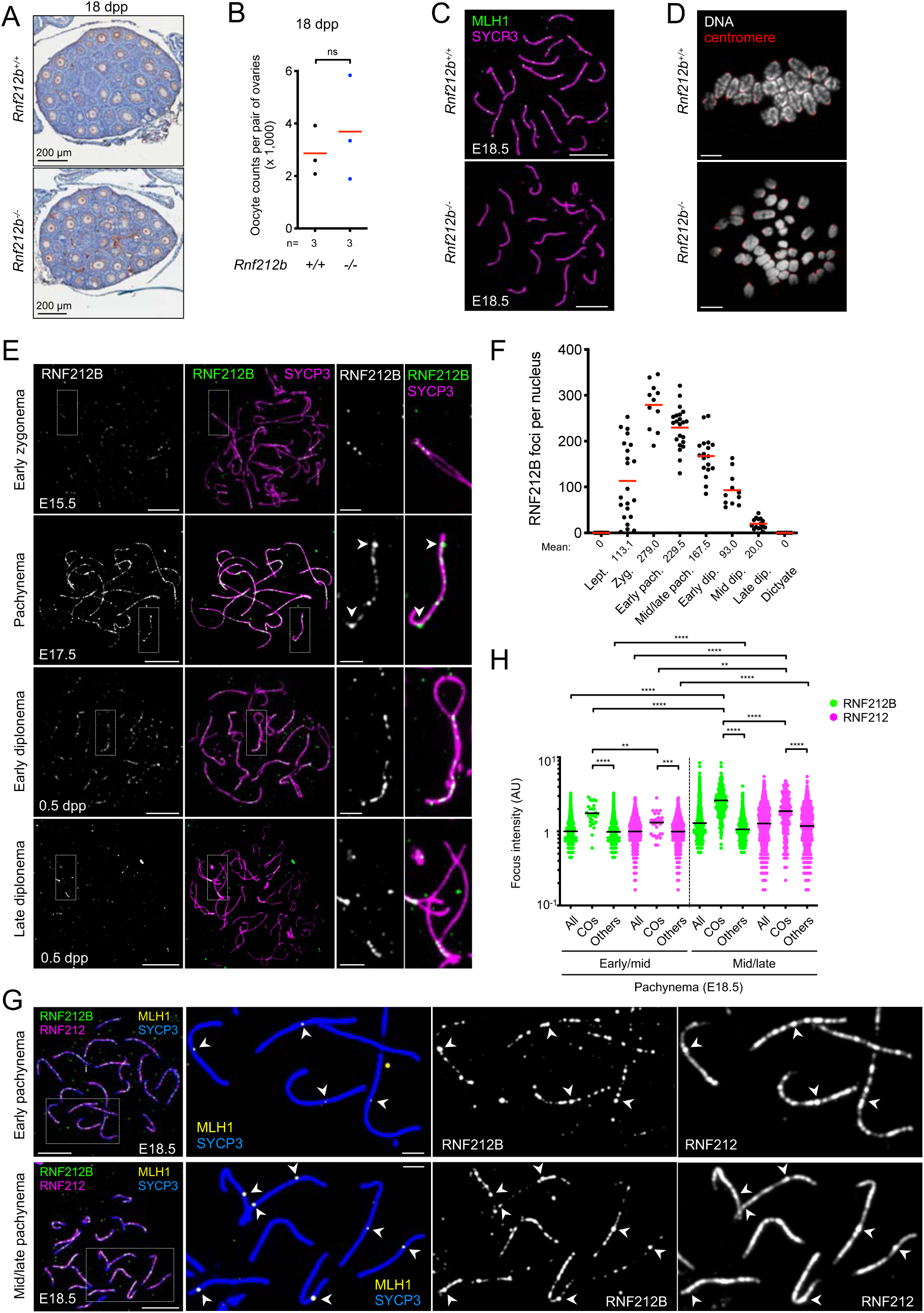
RNF212B is required for crossing over and localizes to the synaptonemal complexes and recombination sites in prophase I oocytes. (A) Ovary sections from wild-type and *Rnf212b^-/-^* mutant females at 18 days postpartum (dpp) immunostained for p63 and counterstained with hematoxylin. (B) Oocyte counts at 18 dpp. Red bars indicate means. ns, not significant (*p* > 0.05, two-tailed *t* test). n, numbers of mice analyzed. (C) MLH1 foci are absent in *Rnf212b^-/-^* oocytes. Pachytene-stage oocyte nuclei from wild-type and *Rnf212b^-/-^* ovaries at embryonic day 18.5 immunostained for SYCP3 and MLH1. Scale bars, 10 μm. (D) Chiasmata are diminished in *Rnf212b^-/-^* oocytes. Metaphase-I oocytes from ovaries of ≥2-month-old wild-type and *Rnf212b^-/-^* females stained with DAPI and immunostained for CREST (centromeres). Scale bars, 10 μm. (E) RNF212B localization at successive prophase-I stages of fetal oocyte nuclei from embryonic days 15.5 (E15.5) and 17.5 (E17.5), and 0.5 day postpartum (0.5 dpp). Surface-spread prophase-I oocyte nuclei were immunostained for SYCP3 and RNF212B. Arrowheads indicate large, differentiated RNF212B foci. The magnified images show representative chromosomal regions. Scale bars, 10 μm for images of full nuclei and 2 μm for magnified panels. (F) Numbers of RNF212B foci in oocyte nuclei at successive prophase-I stages. Red bars indicate means. Means of focus numbers are indicated below the graph. Numbers of nuclei analyzed in leptonema (at E15.5), zygonema (at E15.5), early pachynema (pachytene nuclei with ≤4 MLH1 foci at E16.5), mid/late pachynema (pachytene nuclei with ≥20 MLH1 foci at E18.5), early diplonema (at 0.5 dpp), mid diplonema (at 0.5 dpp), late diplonema (at 0.5 dpp) and dictyate stage (at 0.5 dpp) are 21, 22, 11, 22, 18, 11, 15 and 7, respectively. (G) RNF212B colocalization with RNF212 and MLH1 in oocytes. Early pachytene (top) and mid/late pachytene (bottom) oocytes from E18.5 ovaries immunostained for SYCP3, RNF212B, RNF212 and MLH1. Arrowheads indicate crossover sites. The magnified images show representative chromosomal regions. Scale bars, 10 μm for images of full nuclei and 2 μm for magnified panels. (H) Quantification of focus intensities for RNF212B and RNF212. Black bars indicate means. \*\**p* ≤ 0.01; \*\*\**p* ≤ 0.001; \*\*\*\**p* ≤ 0.0001 for two-tailed Mann-Whitney tests. 7 early/mid pachytene (with ≤10 MLH1 foci) and 9 mid/late pachytene (with ≥20 MLH1 foci) nuclei from E18.5 ovaries were analyzed. Total numbers of foci analyzed: 28 crossover RNF212B foci, 1,375 other RNF212B foci, 30 crossover RNF212 foci and 1,596 other RNF212 foci in early/mid pachynema; 250 crossover RNF212B foci, 1,468 other RNF212B foci, 245 crossover RNF212 foci and 1,580 other RNF212 foci. All, all foci; COs, crossover foci colocalized with MLH1 foci; Others, other foci that don’t colocalize with MLH1 foci.

### Sexually dimorphic chromosomal localization of RNF212B, RNF212, and HEI10

<u>RNF212B</u>: In fetal oocytes, the initial immunostaining pattern of RNF212B during zygonema was similar to that observed in spermatocytes, with foci localized specifically to regions of homolog synapsis (**Figure 6E,F**). Focus numbers also peaked in early pachynema (defined as pachytene-stage oocytes with ≤4 MLH1 foci from embryonic day E16.5 ovaries) but were 1.6-fold greater than in spermatocytes at the same stage (**Figure 6F**; 279.0 ± 50.4 foci per oocyte nucleus, *n =* 11, versus 175.9 ± 15.2 foci per spermatocyte, *n =* 27; mean ± S.D.). Also distinct from spermatocytes, high numbers of RNF212B foci persisted along SCs throughout pachynema both during and after the emergence of amplified foci at designated crossover sites. Indeed, even after a full complement of crossover sites had matured in mid/late pachytene oocytes (with ≥20 MLH1 foci, **Figure 6F**), RNF212B foci averaged 229.5 ± 45.0 per nucleus (from E18.5 ovaries; mean ± SD; *n =* 22; **Figure 6F**). Furthermore, RNF212B remained on synapsed regions throughout diplonema, as homologs desynapsed, and disappeared only as oocytes entered the dictyate stage (**Figure 6E,F**).

Given the distinct dynamics of RNF212B in oocytes, we also analyzed the timing and degree of amplification of RNF212B and RNF212 at crossover sites (**Figure 6G,H**). In early/mid pachytene oocytes (with ≤10 MLH1 foci from E18.5 ovaries), although crossover-associated RNF212B foci were on average 1.8-fold brighter than other (noncrossover) RNF212B foci in the same nuclei (1.76 ± 0.58 for 28 crossover foci versus 0.98 ± 0.40 for 1,375 other foci from 7 nuclei; means ± SDs; *p* < 0.0001, Mann-Whitney test; **Figure 6H**), their intensity distributions overlapped; indeed, we readily observed individual SCs with crossover and non-crossover RNF212B foci of similar intensities (**Figure 6G**, upper panels).

By mid/late-pachynema (oocytes with ≥20 MLH1 foci from E18.5 ovaries), crossover-associated RNF212B foci were now 2.5-fold brighter than other RNF212B foci, suggesting continued amplification after initial designation (2.63 ± 1.13 for 250 crossover foci versus 1.07 ± 0.43 for 1,468 other foci from 9 nuclei; means ± SDs; *p* < 0.0001, Mann-Whitney test; **Figure 6H**). Thus, in oocytes, growth/amplification of RNF212B may be coincident with crossover-site maturation marked by the emergence of MLH1 foci. This contrasts spermatocytes, in which amplified RNF212B foci clearly precede the appearance of MLH1 (**Figure 3**). Again, while crossover associated RNF212B foci were brighter than other foci at the population level, they were not always the brightest foci along the same SC (**Figure 6G**, lower panels). Moreover, a consistent pattern of RNF212B foci at and surrounding designated crossover sites was not observed: some crossover foci were adjacent to other similarly bright foci, others were embedded in a domain of fainter foci, and others by gaps of diminished RNF212B staining (**Figure 6G**). Notably, the intensities of other (noncrossover) RNF212B foci and the total signal intensity of RNF212B per oocyte nucleus were largely unchanged throughout pachynema (**Figure 6H** and **S13A**), in sharp contrast to the loss of RNF212B foci from non-crossover sites in spermatocytes (**Figures 2, 3** and **S7**).

<u>RNF212</u>: Abundant general staining of RNF212 was also retained throughout pachynema in oocytes, but amplification of RNF212 at crossover sites appeared less pronounced than that of RNF212B (**Figure 6G**). Crossover-associated RNF212 foci in early/mid pachynema were only 1.3-fold brighter than other foci (1.32 ± 0.47 for 30 crossover foci versus 0.99 ± 0.45 for 1.596 other foci from 7 nuclei; means ± SDs; *p* = 0.0001, Mann-Whitney test). By mid/late pachynema, crossover-associated RNF212 were now 1.6-fold brighter than other foci in the same nuclei (1.87 ± 1.04 for 245 crossover foci versus 1.19 ± 0.75 for 1,580 other foci form 9 nuclei; means ± SDs; *p* < 0.0001, Mann-Whitney test) and 1.9-fold brighter than those in early/mid pachytene consistent with continued amplification (**Figure 6H**). Intriguingly, the intensity of RNF212 at noncrossover sites slightly increased in mid/late pachynema nuclei suggesting that general loading of RNF212 along SCs may continue during pachytene (0.99 ± 0.45 to 1.19 ± 0.75 from early/mid to mid/late pachynema; means ± SDs; *p* < 0.0001, Mann-Whitney test; **Figure 6H**). Moreover, total signal intensity of RNF212 per oocyte nucleus was largely unchanged between early/mid to mid/late pachynema, as seen for RNF212B (**Figure S13A**).

<u>HEI10</u>: Chromosomal localization of HEI10 was also sexually dimorphic. Unlike spermatocytes, prominent HEI10 foci were already apparent in zygotene-stage oocytes, specifically at regions of synapsis (**Figure S13B**). Focus numbers increased as synapsis ensued, peaking in early pachynema (defined as pachytene-stage oocytes with ≤4 MLH1 foci) with 123.9 ± 28.7 foci per nucleus (mean ± SD; *n =* 18 nuclei; **Figure 7A,D**). At this stage, HEI10 foci were unevenly spaced and highly variable in size and intensity; with configurations suggestive of growth, shrinkage, and/or fusion of foci as might be predicted by coarsening models (**Figure 7A,F** and **S13B,D**). Focus numbers then reduced as pachynema progressed, showing a largely crossover-specific pattern by mid pachynema (defined as pachytene-stage oocytes with ≥20 MLH1 foci from E17.5 ovaries; **Figure 7 B–E**). At this stage, 86.2 ± 9.6 % of HEI10 foci colocalized with MLH1 foci and 98.0 ± 2.7 % of MLH1 foci colocalized with HEI10 foci (means ± SDs; *n =* 14 nuclei).

**Figure 7.**
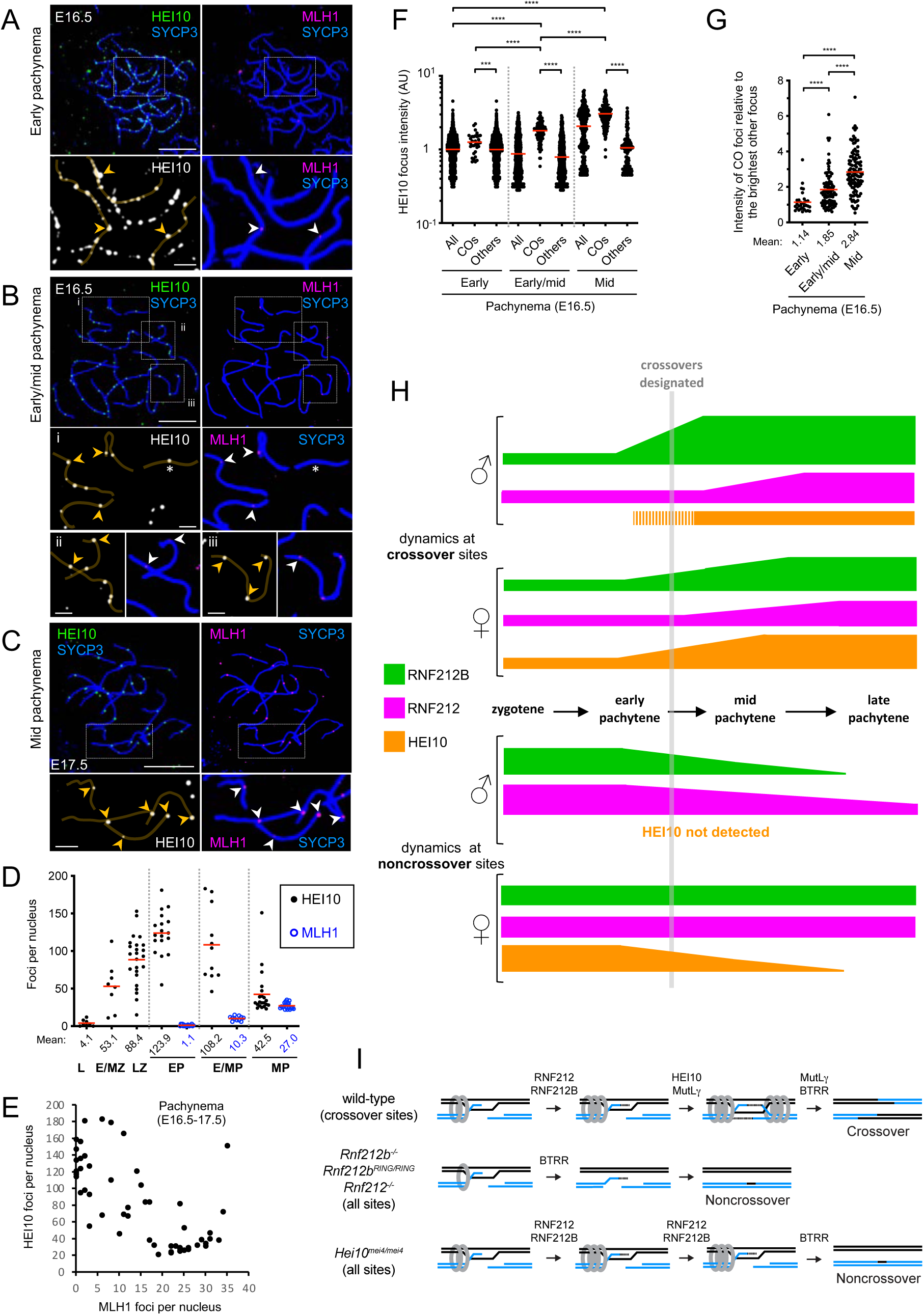
Dynamic chromosomal localization of HEI10 in oocytes. (A-C) HEI10 localization at successive prophase-I stages of fetal oocyte nuclei from embryonic days 16.5 (E16.5) and 17.5 (E17.5). Surface-spread prophase-I oocyte nuclei in early pachynema (A), early/mid pachynema (B), and mid pachynema (C) were immunostained for SYCP3, HEI10 and MLH1. Arrowheads indicate crossover foci marked by MLH1. The asterisk indicates an SC with an isolated bright HEI10 focus but without an MLH focus. The magnified images show representative chromosomal regions. Scale bars, 10 μm for images of full nuclei and 2 μm for magnified panels. (D) Numbers of HEI10 and MLH1 foci in oocyte nuclei at successive prophase-I stages. Red bars indicate means. Means of focus numbers are indicated below the graph. Numbers of nuclei analyzed in leptonema (L; at E16.5), early and mid zygonema (E/MZ; at E16.5), late zygonema (LZ; at E16.6), early pachynema (pachytene nuclei with ≤4 MLH1 foci at E16.5), early-to-mid pachynema (E/MP; pachytene nuclei with 5-15 MLH1 foci at E16.5), and mid pachynema (MP; pachytene nuclei with ≥20 MLH1 foci at E16.5 and E17.5), are 8, 8, 24, 18, 11 and 21, respectively. (E) Quantification of HEI10 and MLH1 foci in pachytene oocytes from E16.5 and E17.5 ovaries. 56 pachytene nuclei were randomly selected and analyzed. (F) Quantification of HEI10 focus intensities. Red bars indicate means. \*\*\**p* ≤ 0.001; \*\*\*\**p* ≤ 0.0001 for two-tailed Mann-Whitney tests. 26 early pachytene (with ≤4 MLH1 foci), 9 early/mid pachytene (with 5-15 MLH1 foci) and 6 mid pachytene (with ≥20 MLH1 foci) nuclei from E16.5 ovaries were analyzed. Total numbers of foci analyzed: 33 crossover foci and 3,817 other foci in early pachynema; 96 crossover foci and 1,089 other foci in early/mid pachynema; 186 crossover foci and 180 other foci. All, all foci; COs, crossover foci colocalized with MLH1 foci; Others, other foci that don’t colocalize with MLH1 foci. (G) Per chromosome analysis for HEI10 focus intensity. Intensity of each crossover-associated HEI10 focus relative to that of the brightest other (non-crossover) focus along the same chromosome. Red bars indicate means. Mean intensities are indicated below the graph. Detailed representation of per chromosome analysis is shown in **Figure S13B**. \*\*\*\**p* ≤ 0.0001 for two-tailed Mann-Whitney tests. 31, 92, and 111 crossover foci from 14 early pachytene, 9 early/mid pachytene, and 6 mid pachytene nuclei were analyzed, respectively. (H) Summary of localization dynamics of RNF212B, RNF212, and HEI10 at designated crossover sites and noncrossover sites in spermatocytes and oocytes. See text for details. (I) Models of crossover maturation and defects in *Rnf212b, Rnf212* and *Hei10* mutants. MutSγ complexes (grey rings) bind nascent Holliday junctions and convert into sliding clamps that embrace the interacting duplexes ^28^. At designated crossover sites, stabilization of MutSγ (and other ZMMs) by RNF212 and RNF212B stabilize single-end invasions. HEI10 and an endonuclease-independent function of MutLγ^58^ then facilitate second-end capture to form dHJs. MutLγ-catalyzed DNA incision and the Bloom complex (BLM-TOPIIIα-RMI1-RMI2; BTRR) then mediate crossover-specific dHJ resolution ^29^. MutSγ is destabilized in *Rnf212b^-/-^, Rnf212b^RING/RING^* and *Rnf212^-/-^* mutants exposing nascent intermediates to unwinding by the Bloom complex resulting in noncrossovers. In the *Hei10^mei4/mei4^* mutant, RNF212, RNF212B, and MutSγ persist at all sites throughout pachytene, and crossover sites fail to differentiate. DSBs are repaired as noncrossovers only as homologs desynapse and RNF212 and RNF212B dissociate.

Crossover-associated HEI10 foci in early-to-mid pachynema (pachytene-stage oocytes with 5-15 MLH1 foci from E16.5 ovaries) were on average 2.3-fold brighter than other foci in the same nuclei (1.80 ± 0.53 for 96 crossover foci versus 0.79 ± 0.48 for 1,089 other foci from 9 nuclei; means ± SDs; *p* < 0.0001, Mann-Whitney test; **Figure 7B,F**). Even very faint emergent MLH1 foci in early pachynema (pachytene-stage oocytes with ≤4 MLH1 foci from E16.5 ovaries) tended to be associated with slightly larger HEI10 foci suggesting that amplification accompanies crossover designation (1.26 ± 0.39 for 33 crossover foci versus 1.00 ± 0.45 for 3,817 other foci from 26 nuclei; means ± SDs; *p* = 0.0003, Mann-Whitney test; **Figure 7A,F**). However, MLH1 foci emerged when HEI10 foci still outnumbered crossovers by more than 3:1, i.e. before HEI10 *per se* had attained a crossover-specific distribution. Moreover, while crossover-associated HEI10 foci were on average 1.9-fold brighter relative to the brightest other focus along the same chromosome in early/mid pachynema (**Figure 7G**), similarly bright HEI10 foci were often observed along the same SC (**Figure 7B**, magnified panels, and **Figure S13B,C**). After mid pachynema (oocytes with ≥20 MLH1 foci from E16.5 ovaries), crossover-associated HEI10 foci were 3.1-fold brighter than the foci detected in early pachynema and 1.7-fold brighter than the crossover-specific foci detected in early/mid pachynema (3.06 ± 1.00 for 186 crossover foci from 6 nuclei; mean ± SD; **Figure 7F**), suggesting continued amplification of HEI10 after crossover designation. The intensities of other remaining (noncrossover) HEI10 foci in mid pachynema were comparable to those of the foci detected in early pachynema, though these foci were eventually lost (**Figure 7D-F**).

Together, these data reveal profound sexual dimorphism in the localization dynamics of RNF212B, RNF212, and HEI10. Importantly, in oocytes, designation of crossover sites does not involve general depletion of RNF212B and RNF212 from noncrossover sites; and amplification may be a coincident or downstream event. In contrast, as crossover associated HEI10 foci amplify, foci at other sites progressively diminish in number. While the timing of HEI10 amplification may be consistent with a role in crossover-site designation, crossover sites are maturing well before a crossover-specific pattern of HEI10 is attained.

## DISCUSSION

### COR Patterning vis-a-vis Crossover Patterning

An elementary coarsening model, that can explain crossover patterning in Arabidopsis, posits that a finite amount of HEI10 protein (fixed amount per unit length) diffuses along SCs and becomes progressively absorbed into one or a few amplified foci which then designate which recombination sites will mature into crossovers ^35, 36^. Our data in mouse are hard to reconcile with this model, revealing a more complex scenario in which pattering of a given COR protein may precede, accompany, or follow the designation of crossover sites; can occur with or without depletion of CORs from non-crossover sites; and shows striking sexual dimorphism (summarized in **Figure 7H**).

### Global and local regulation of recombination by RNF212B, RNF212, and HEI10

What might be the function(s) of the observed dynamics of RNF212, RNF212B and HEI10? Our analysis emphasizes the interdependencies and distinctions between the three mammalian CORs and their global and local functions that help coordinate key events of meiotic prophase I. Most evident is regulating the progression of recombination occurring in the context of the SC. Following initial DNA strand-exchange and homolog pairing, RNF212-RNF212B associates with nascent SCs and acts to stabilize ZMM factors and pause the progression of recombination. In this way, RNF212-RNF212B may act to protect nascent recombination intermediates from being prematurely dissociated by the Bloom complex, which is inferred to mediate the default noncrossover outcome of recombination via synthesis-dependent strand annealing ^31–34, 53^. This early function of RNF212-RNF212B appears to promote and stabilize synapsis, possibly acting as a kind of proofreading mechanism that selectively reinforces SC assembled at recombination sites, i.e. between homologous chromosomes. As prophase-I progresses, connection of homologs by recombinational interactions is superseded by the SC. RNF212-RNF212B could mediate this hand-off by ensuring that recombinational connections are not resolved until mature SC has formed.

Pausing the progression of SC-associated recombination events may also be a prerequisite for crossover/noncrossover differentiation and/or to allow time for crossover sites to mature. Importantly, RNF212-RNF212B renders the progression of recombination dependent on HEI10 (**Figure 4**) and additional pro-crossover factors including kinase CDK2, and the CNTD1-PRR19 complex ^16, 17, 45, 54, 55^. In the absence of HEI10, foci of RNF212-RNF212B and ZMM proteins remain abundant and undifferentiated along SCs, and recombination continues to be stalled throughout pachytene, with DSB repair being completed only as the SC central region disassembles along with RNF212-RNF212B.

At designated crossover sites, local protection of intermediates via RNF212-RNF212B may enable crossover-specific events including dHJ formation and recruitment of the crossover-specific resolution machinery, organized around the MutLγ endonuclease. Local stabilization of SC via RNF212-RNF212B may also explain why crossover sites are the last sites to desynapse during diplotene ^56^. We have suggested that patches of SC retained at crossover sites may help coordinate the DNA events of crossing over with exchange of the underlying chromosome axes to form chiasmata.

### Distinct and interdependent functions of RNF212, RNF212B and HEI10

Our ability to discern potentially distinct functions of RNF212 and RNF212B is compromised by their interdependence for protein stability, i.e. *Rnf212* and *Rnf212b* single mutant phenotypes likely reflect diminished function of both proteins. However, localization dynamics of RNF212 and RNF212B reveal divergent behaviors of the two proteins that point to distinct function and regulation.

Notably, in early pachytene spermatocytes, RNF212 foci outnumber RNF212B and up to 40% don’t detectably colocalize. Also, the amplification of crossover specific RNF212B foci was much stronger than for RNF212 and occurred earlier, with the suggestion that differentiation may occur via distinct processes, e.g. redistribution to crossover sites for RNF212B, versus the loss from noncrossover sites for RNF212 (or via different proportions of redistribution and loss). Thus, although RNF212 and RNF212B interact, these observations argue against the existence of an obligate heterocomplex of fixed stoichiometry and suggest that the activity of RNF212B may be modulated by switching binding partners as it accumulates at designated crossover sites. For example, RNF212B-RNF212B self-interaction could become prominent, or RNF212B could partner with HEI10 in order to locally accumulate and modulate E3 ligase activities at designated crossover sites. Consistent with the latter possibility, HEI10-RNF212B interaction is detected by Y2H, and differentiation of RNF212B foci is both dependent on HEI10 and temporally indistinguishable from its appearance at prospective crossover sites (**Figures 4** and **S8**).

Localization dynamics in mutant backgrounds also point to distinct functions and regulation of the three CORs. For example, RNF212 localization along SCs remains robust in the absence of SPO11-catalyzed DSBs while localization of RNF212B is much less efficient, suggesting that DSB signaling may stabilize RNF212B. In this respect, RNF212B may be like budding yeast Zip3, which requires phosphorylation at S/T-Q consensus sites for the DNA-damage response (DDR) kinases Mec1^ATR^/Tel1^ATM^ for normal localization ^57^. Further distinctions are seen in the absence of synapsis, with RNF212 showing a robust ability to localize to recombination sites, while RNF212B localization was much weaker. Thus, inputs from both DSBs and synapsis appear to stabilize RNF212B and thereby coordinate its activity in space and time, while RNF212 might act more as an anchor for localization to SCs and recombination sites. We previously showed that HEI10 foci in spermatocytes are largely dependent on both DSBs and synapsis ^16^, consistent with our inference that HEI10 foci assemble only as crossover sites differentiate, dependent on RNF212-RNF212B.

Defective crossover maturation in the *Mlh3* mutant impacts the dynamics of RNF212, RNF212B and HEI10 to varying degrees. Although crossover-specific patterning of RNF212-RNF212B foci still occurs in *Mlh3* spermatocytes, numerous small foci of RNF212, and to a lesser extent RNF212B, also persist along SCs implying that a MutLγ-dependent signal associated with the maturation of crossover sites enhances the general loss of RNF212-RNF212B from noncrossover sites. We previously showed that HEI10 patterning is severely perturbed in *Mlh3* mutant spermatocytes, with abnormally high numbers of foci (∼90 per nucleus) forming much earlier than normal (in zygonema) and then persisting at recombination sites until chromosomes desynapse ^16^. Thus, although HEI10 is required for crossover patterning of RNF212-RNF212B foci, this function does not require crossover-specific patterning of HEI10 itself, i.e. crossover-specific localization of HEI10 appears to be downstream of initial crossover designation. Moreover, crossover-specific patterning of HEI10 requires an upstream maturation step that is dependent on the MutLγ complex, likely the dHJ formation function described by Premkumar et al. ^58^. Consistently, in oocytes, MLH1 foci can be detected before HEI10 foci have attained a crossover-specific pattern (**Figure 7A** and **S13BC**).

Collectively, these data suggest that RNF212, RNF212B and HEI10 may function as apical effectors that coordinate meiotic prophase by integrating signals from synapsis, DSB repair, cell cycle kinases, and maturating crossover sites (summarized in **Figure 7**).

### Sexual dimorphism

Patterning of all three CORs along SCs shows striking sexual dimorphism. In spermatocytes, complete differentiation of RNF212B results in only large crossover-specific foci; while in oocytes, numerous foci persist throughout pachytene even after amplified crossover-specific foci have formed and crossover sites have matured (marked by the appearance of MLH1 foci; **Figures 2A,H**, and **6E,G**). RNF212 dynamics are similarly dimorphic. Thus, complete differentiation of RNF212B or RNF212 is not a prerequisite for the patterning and maturation of crossover sites. Possibly, accumulation of a threshold level of RNF212B at a given recombination site is sufficient for it to mature into a crossover. However, weak amplification and incomplete differentiation in females could render RNF212B crossover foci less stable and potentially reversible, possibly accounting for the inefficiency of crossover maturation characterized in human oocytes that can result in unconnected (achiasmate) homologs, or homologs connected only by a single telomere-proximal chiasma, configurations that are prone to segregation error ^8, 9^. HEI10 shows striking temporal and spatial dimorphism with foci emerging in mid pachytene spermatocytes already showing a crossover distribution; while in oocytes, numerous foci first form during zygotene and then attain a crossover-specific pattern in pachytene.

What could account for the sexually dimorphic behavior of mammalian CORs? A pertinent difference may be the duration of pachytene, which lasts almost 7 days in mouse spermatocytes^59^ compared to just 1-2 days in oocytes ^60^. Thus, if patterning dynamics are relatively slow, differentiation may remain incomplete in oocytes. Also, oocyte SCs are longer than spermatocytes and CORs appear to be generally more abundant. COR dimorphism might also reflect dimorphism in the DNA-damage checkpoint and gamete quality control. We previously showed that RNF212 is required for oocyte quality control as dictyate oocytes assemble into primordial follicles ^61^, while the primary checkpoint in spermatocytes occurs much earlier, as the sex-body assembles in mid pachytene ^62^.

## SUPPLEMENTAL FIGURES

**Figure S1.**
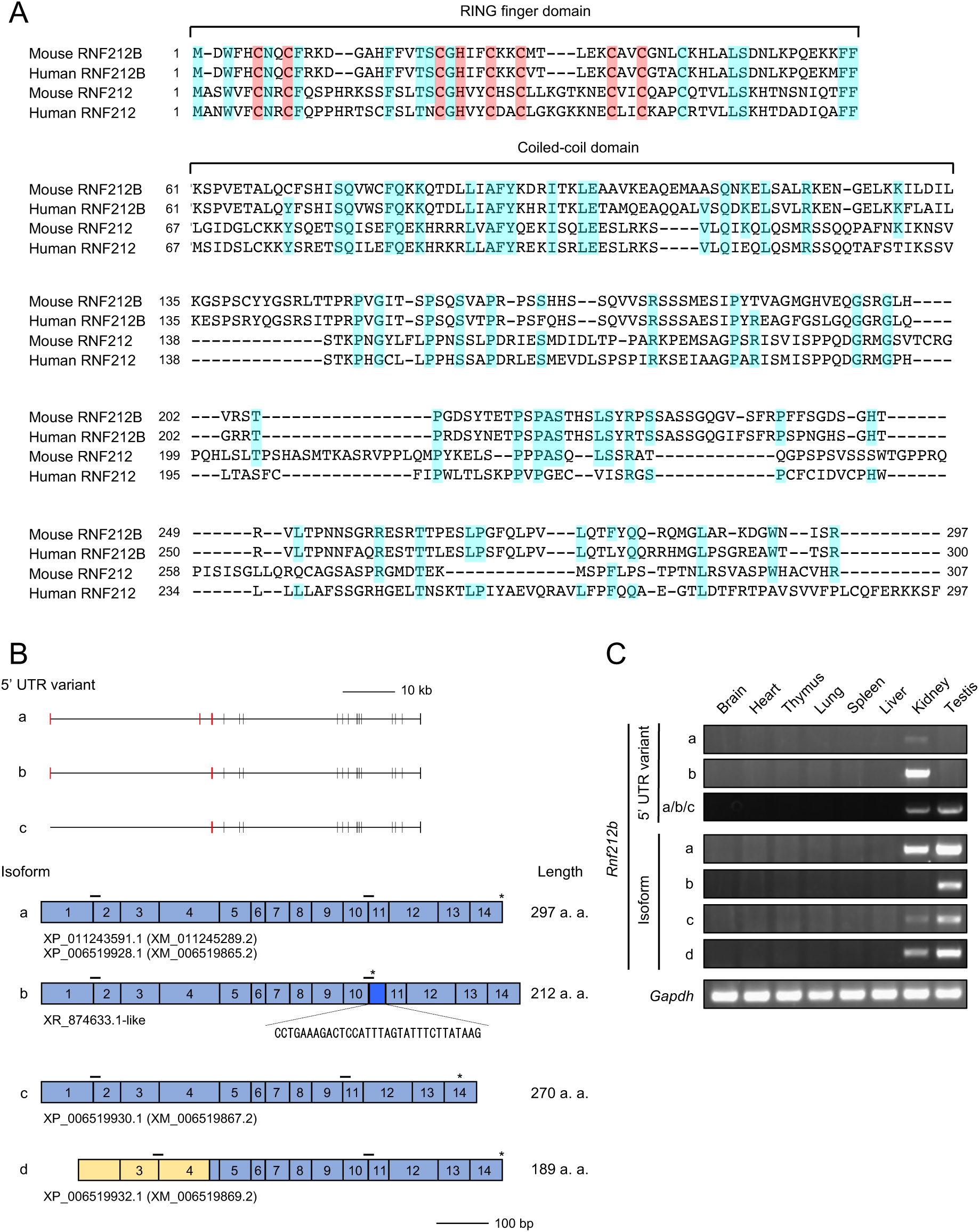
5’ UTR variants and isoforms of mouse *Rnf212b*, related to Figure 1. Protein sequence alignments of mouse and human RNF212B and RNF212 full-length isoforms. Zinc-coordinating cysteine and histidine (pink) residues of RING finger domain and other identical residues (blue) are highlighted. (A) Schematic of 5’ UTR variants and isoforms of mouse *Rnf212b*. Top, red and black bars represent 5’ untranslated exons (5’ UTRs) and coding exons, respectively. Two 5’ UTR variants were identified in the NCBI database, *variants a* and *b*, which differ by a deletion of the second untranslated exon in *variant b*. A third *variant c* was identified by 5’ RACE from testes and comprises ∼1/3 of the third untranslated exon of *variant a* followed by the 14 coding exons. Bottom, yellow and blue boxes represent 5’ UTRs and coding exons, respectively. Four different coding isoforms were identified in the NCBI database. For simplicity, 5’ UTRs of isoforms a-c are not shown. Isoform a encodes the full-length RNF212B protein of 297 amino acids; isoform b uses an alternative splice acceptor site, creating a premature stop codon after exon 10 that is predicted to produce a protein lacking 86 C-terminal amino acids; isoform c uses an alternative splice donor site to precisely skip exon 10, which corresponds to 16 amino acids of the disordered serine-rich C-terminal tail, resulting in a predicted protein of 270 amino acids; and isoform d lacks the first two exons present in other isoforms and uses an extended exon 3 and most of exon 4 as a 5’ UTR, to encode a predicted protein of 189 amino acids that lacks the N-terminal RING finger domain. Asterisks indicate positions of stop codons. Horizontal bars represent positions of primers used to detect each isoform by RT-PCR in (C). Annotated or related variants and isoforms in the NCBI database are also shown. Expression of full-length a and b isoforms in testis was confirmed by *de novo* cloning from testis mRNA (the sequence of cloned isoform b was similar but not identical to XR_874633). (B) Expression of mouse *Rnf212b* 5’ UTR variants and isoforms analyzed by RT-PCR. Total RNA was extracted from indicated tissues of mature male mice and subjected to RT-PCR. *Gapdh* is a loading control.

**Figure S2.**
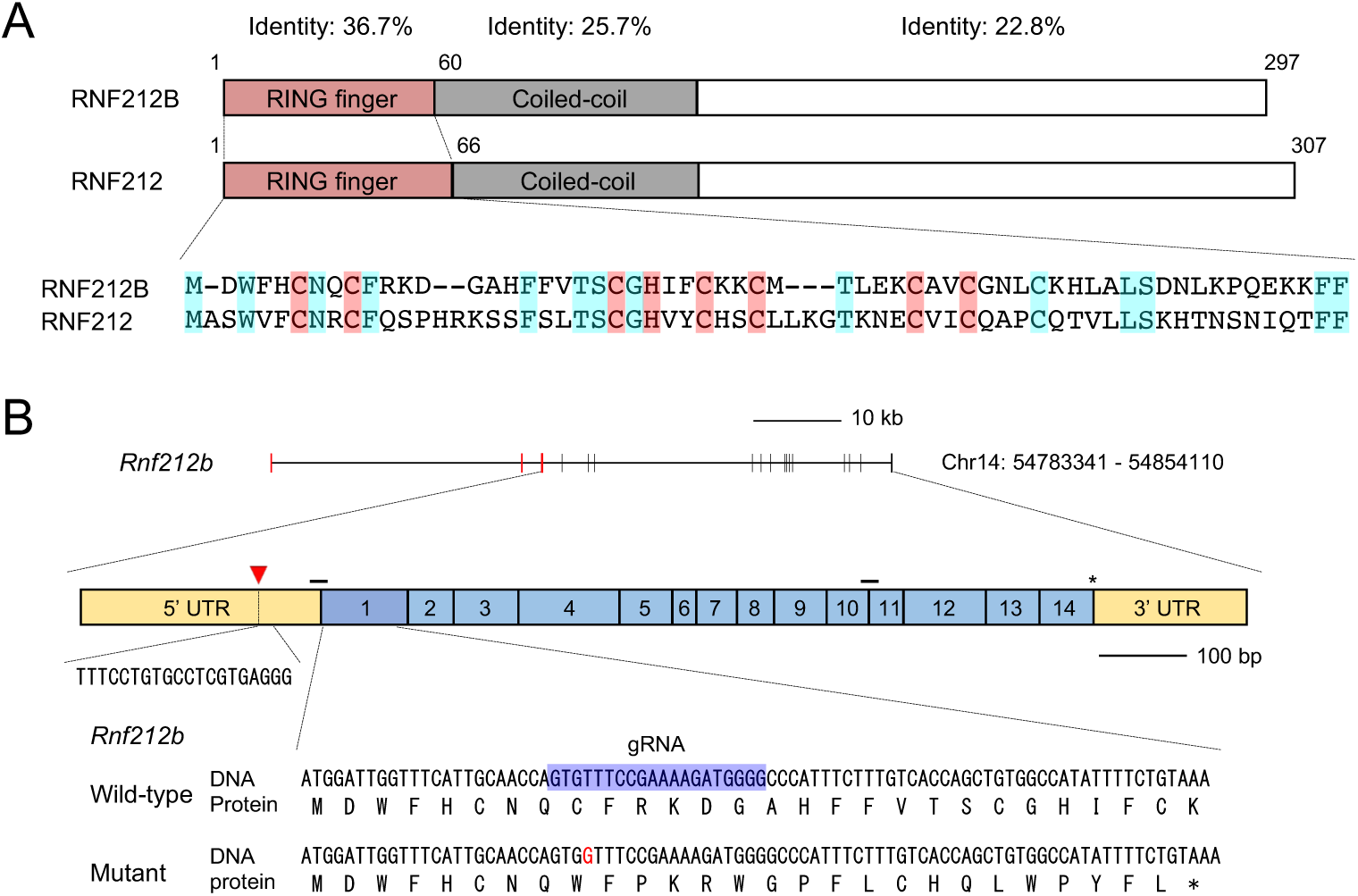
Domain structure of RNF212B and strategy to generate Rnf212b mutant mice by CRISPR/Cas9, related to Figure 1. (A) Domain structures of mouse RNF212B and RNF212. Amino acid sequences of the RING finger domains are shown below. Presumptive zinc-coordinating cysteines and histidines and other identical residues are highlighted in pink and blue, respectively. The domain structure of the full-length RNF212B protein mirrors that of RNF212 with an N-terminal RING finger domain (36.7% amino-acid identity with RNF212), a ∼70 amino acid region of predicted coiled-coil (25.7% identical) and a largely disordered serine-rich C-terminal tail (22.8% identical). (B) Schematic of the mouse *Rnf212b* gene on chromosome 14. Top row: red and black bars represent 5’ untranslated exons (5’ UTRs) and coding exons, respectively. Middle row: yellow and blue boxes represent 5’ UTRs and coding exons, respectively, in the full-length isoform a. The inverted red triangle represents the translation initiation site in testis identified by 5’ RACE. An asterisk represents the stop codon. Horizontal bars represent the positions of primers used to detect 5’ UTR variants a/b/c by RT-PCR in **Figure S1C**. Bottom row: nucleotide and amino-acid sequences at the CRISPR/Cas9 targeted site in exon 1. A single-nucleotide insertion (shown in red) created a frameshift and premature stop codon (asterisk). The guide RNA (gRNA) sequence is highlighted in blue.

**Figure S3.**
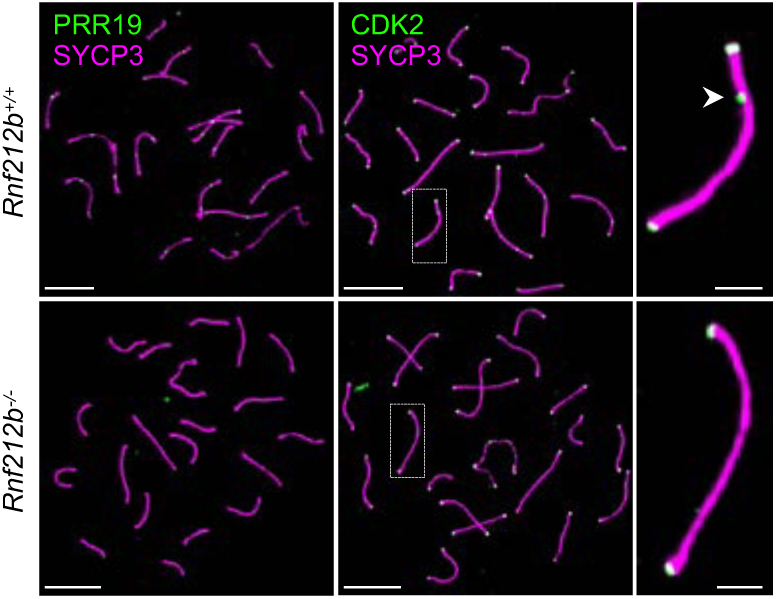
Absence of crossover markers in *Rnf212b^-/-^* mutant mice, related to Figure 1. Crossover-specific immunostaining foci of PRR19 (left) and CDK2 (right) foci were absent from chromosome spreads of *Rnf212b^-/-^* spermatocytes. Mid/late pachytene nuclei from wild-type and *Rnf212b^-/-^* spermatocytes were immunostained for SYCP3 plus PRR19 (left) or CDK2 (right). The magnified images show representative chromosomes from the CDK2 staining. Note that CDK2 also localizes to telomeres, which is not affected by *Rnf212b* mutation. An arrowhead indicates an interstitial CDK2 focus marking a prospective crossover site. Scale bars, 10 μm for images of full nuclei and 2 μm for magnified panels.

**Figure S4.**
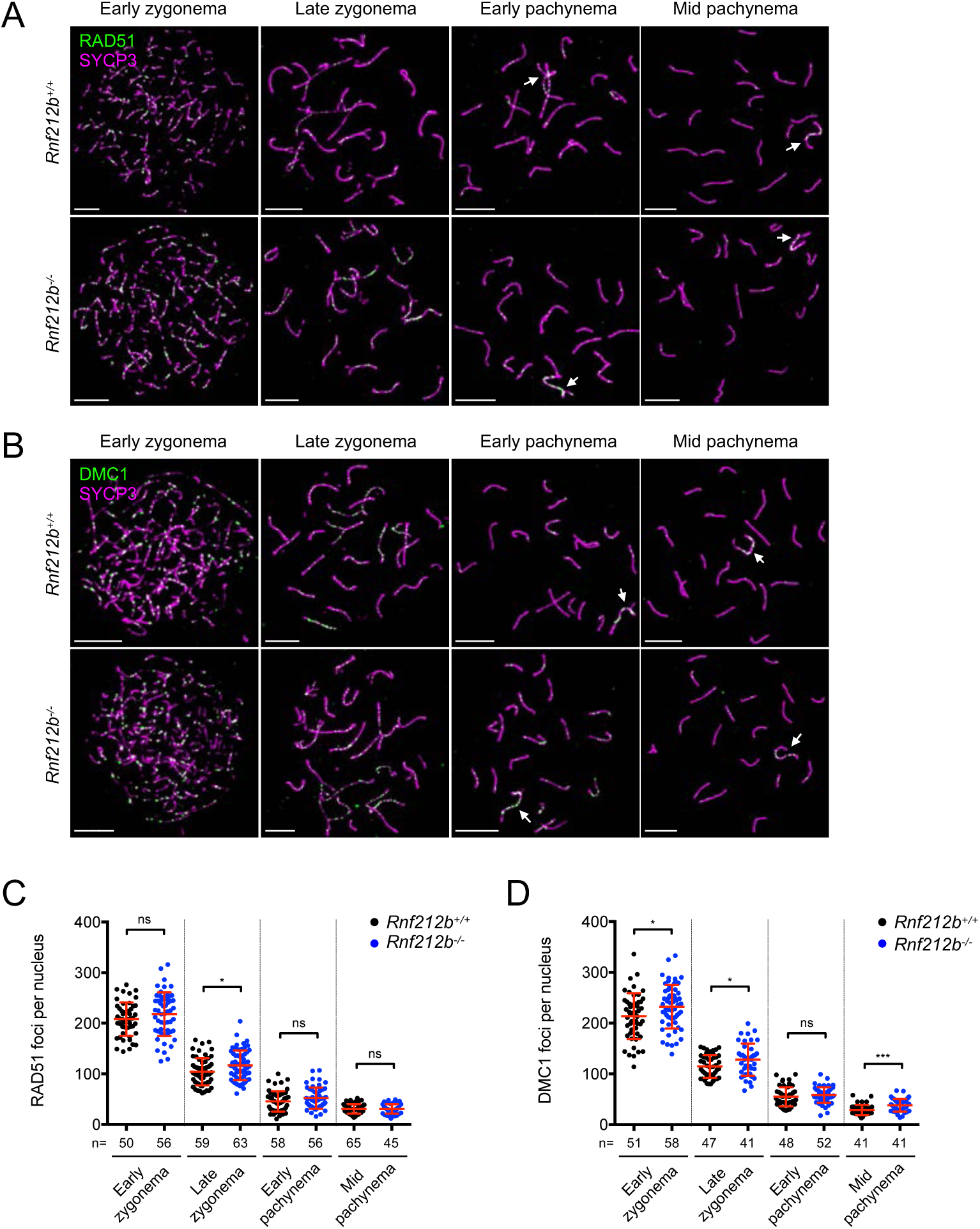
Dynamics of RAD51 and DMC1 throughout zygonema and pachynema, related to Figure 1. (A and B) Representative images of RAD51 (A) and DMC1 (B) immunostainings in wild-type and *Rnf212b^-/-^* spermatocytes nuclei immunostained for SYCP3 plus RAD51 (A) or DMC1 (B) at the indicated stages. Arrows indicate the X-Y chromosomes. Scale bars, 10 μm. (C and D) Focus counts of RAD51 (C) and DMC1 (D). Red bars indicate means ± SDs. ns, not significant (*p* > 0.05); \**p* ≤ 0.05; \*\*\**p* ≤ 0.001 for two-tailed Mann-Whitney tests. Total numbers of nuclei analyzed are indicated below the X axes.

**Figure S5.**
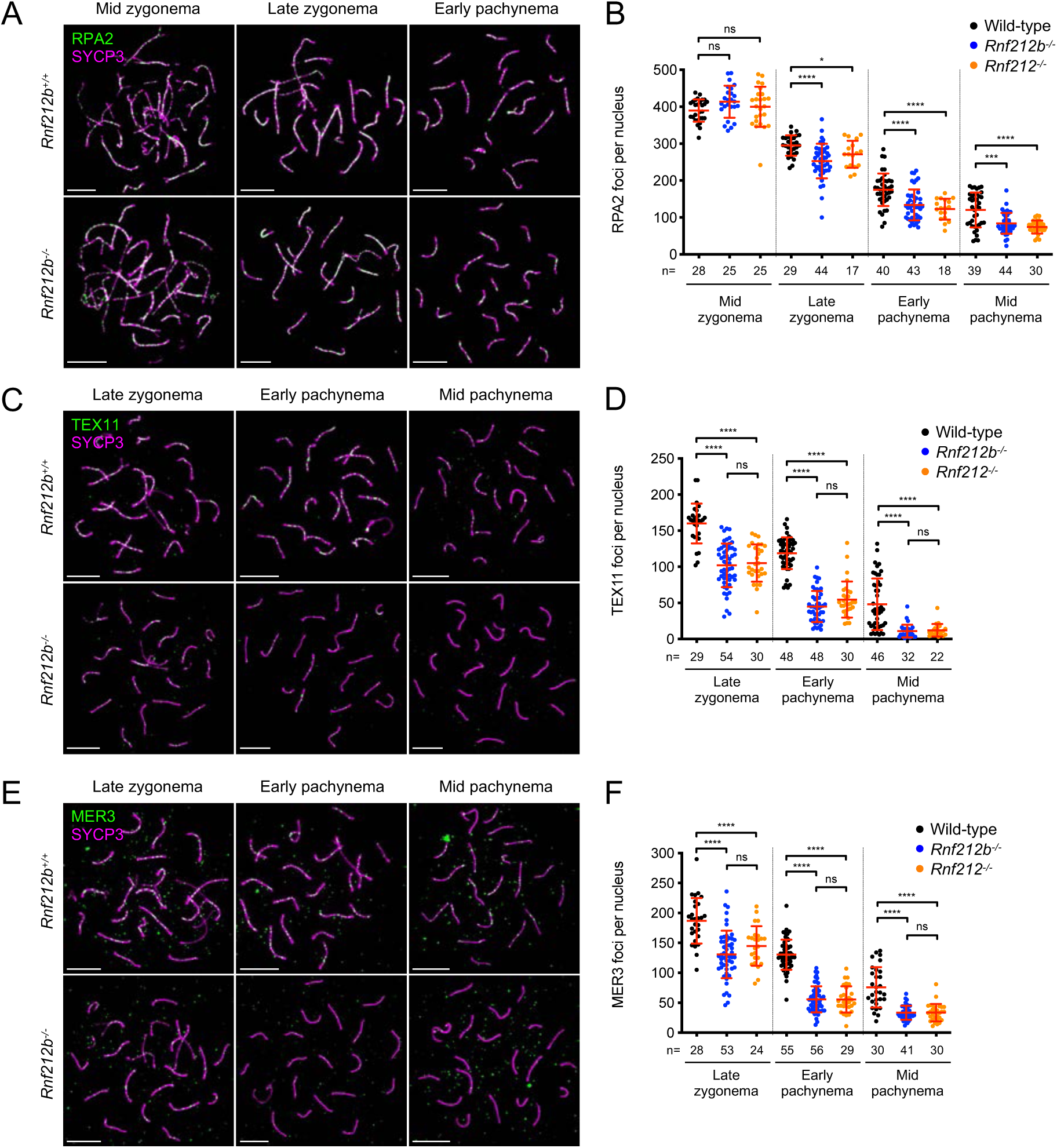
Intermediate steps of recombination are perturbed in *Rnf212b^-/-^* mutant mice, related to Figure 1. (A, C and E) Images of wild-type and *Rnf212b^-/-^* spermatocyte nuclei immunostained for SYCP3 plus RPA2 (A), TEX11 (C) or MER3 (E) at the indicated stages. Scale bars, 10 μm. (B, D and F) Counts of RPA2 (B), TEX11 (D) and MER3 (F) immunostaining foci. Red bars indicate means ± SDs. ns, not significant (*p* > 0.05); \**p* ≤ 0.05; \*\*\**p* ≤ 0.001; \*\*\*\**p* ≤ 0.0001 for two-tailed Mann-Whitney tests. Total numbers of nuclei analyzed are indicated below the X axes.

**Figure S6.**
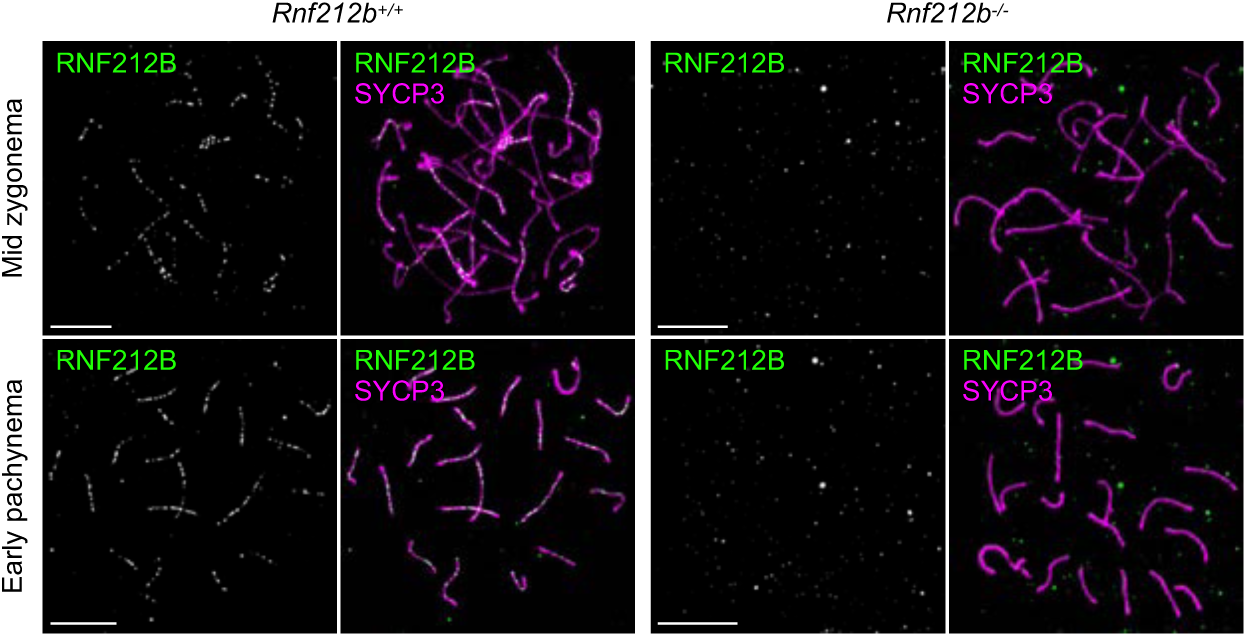
Specificity of anti-RNF212B antibody, related to Figure 2. Wild-type and *Rnf212b^-/-^* spermatocyte nuclei at indicated stages were immunostained for SYCP3 and RNF212B. Scale bars, 10 μm.

**Figure S7.**
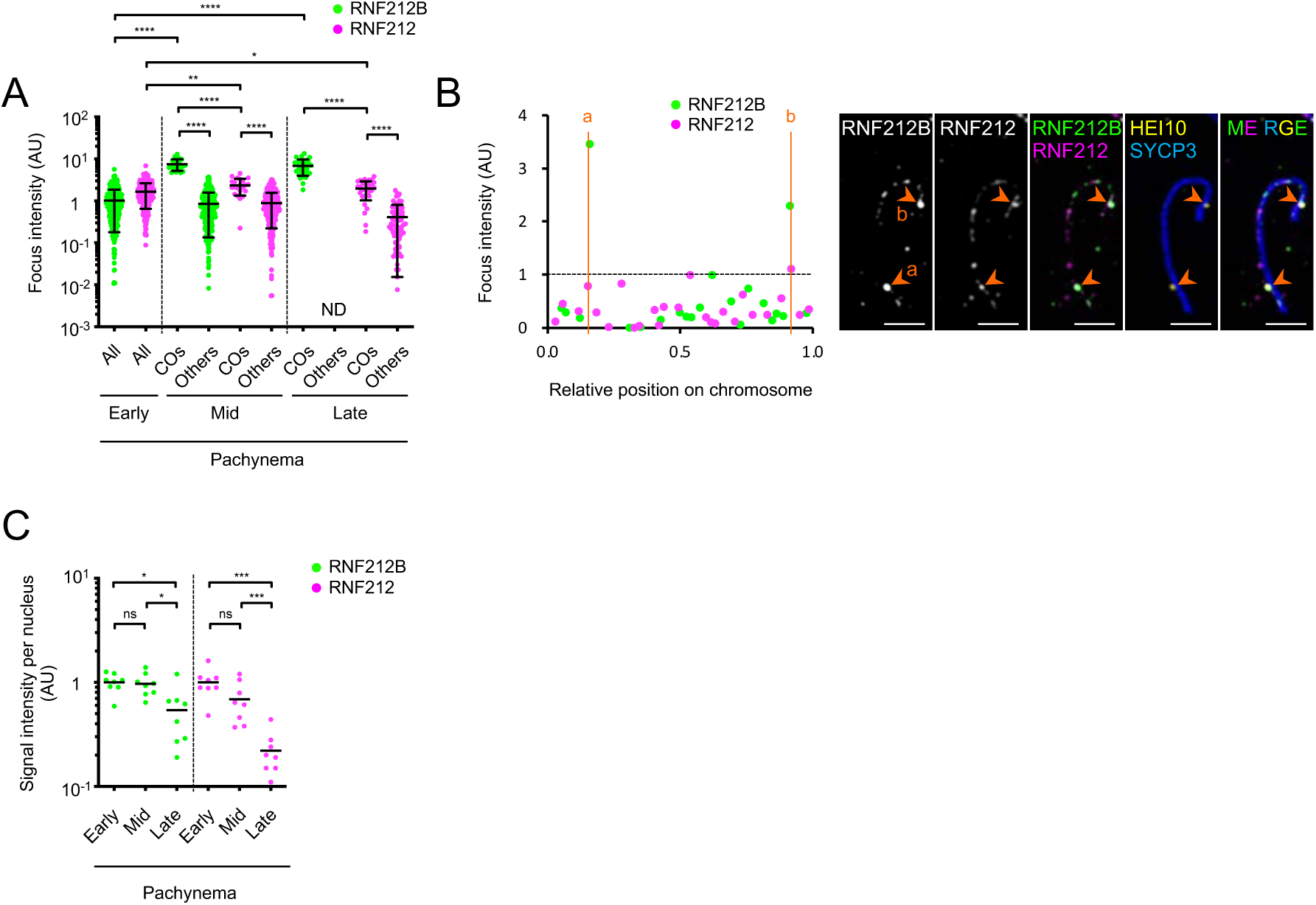
Distinct localization dynamics of RNF212B and RNF212 foci, related to Figure 3. (A) Quantification of focus intensities for RNF212B and RNF212. Each dot represents each focus of RNF212B and RNF212 from the early, mid and late pachytene nuclei (one nucleus per stage) analyzed in Figure 3D. Bars indicate means ± SDs. All, all foci; COs, crossover-specific foci that colocalize with HEI10 foci; Others, foci that don’t colocalize with HEI10 foci (noncrossover foci). ND, not detected. \**p* ≤ 0.05; \*\**p* ≤ 0.01; \*\*\*\**p* ≤ 0.0001 for two-tailed Mann-Whitney tests. (B) Representation of per chromosome analysis of RNF212B and RNF212 foci for a mid-pachytene chromosome (shown in the images in the right-hand panels). Each focus of RNF212B or RNF212 is represented by a dot. The dashed line indicates the signal intensity of the brightest other focus (noncrossover focus) along the same chromosome. Orange vertical lines indicate the positions of HEI10-positive crossover sites (a) and (b). Intensities of foci at these sites relative to that of the brightest other (noncrossover) focus are: (a) 3.5 and (b) 2.3 for RNF212B; and (a) 0.8 and (b) 1.1 for RNF212. (C) Total signal intensity of RNF212B and RNF212 per nucleus in early, mid and late pachytene spermatocytes. Black bars indicate means. ns, not significant (*p* > 0.05); \**p* ≤ 0.05; \*\*\**p* ≤ 0.001 for two-tailed Mann-Whitney tests. Nuclei analyzed were the same as in Figure 3D.

**Figure S8.**
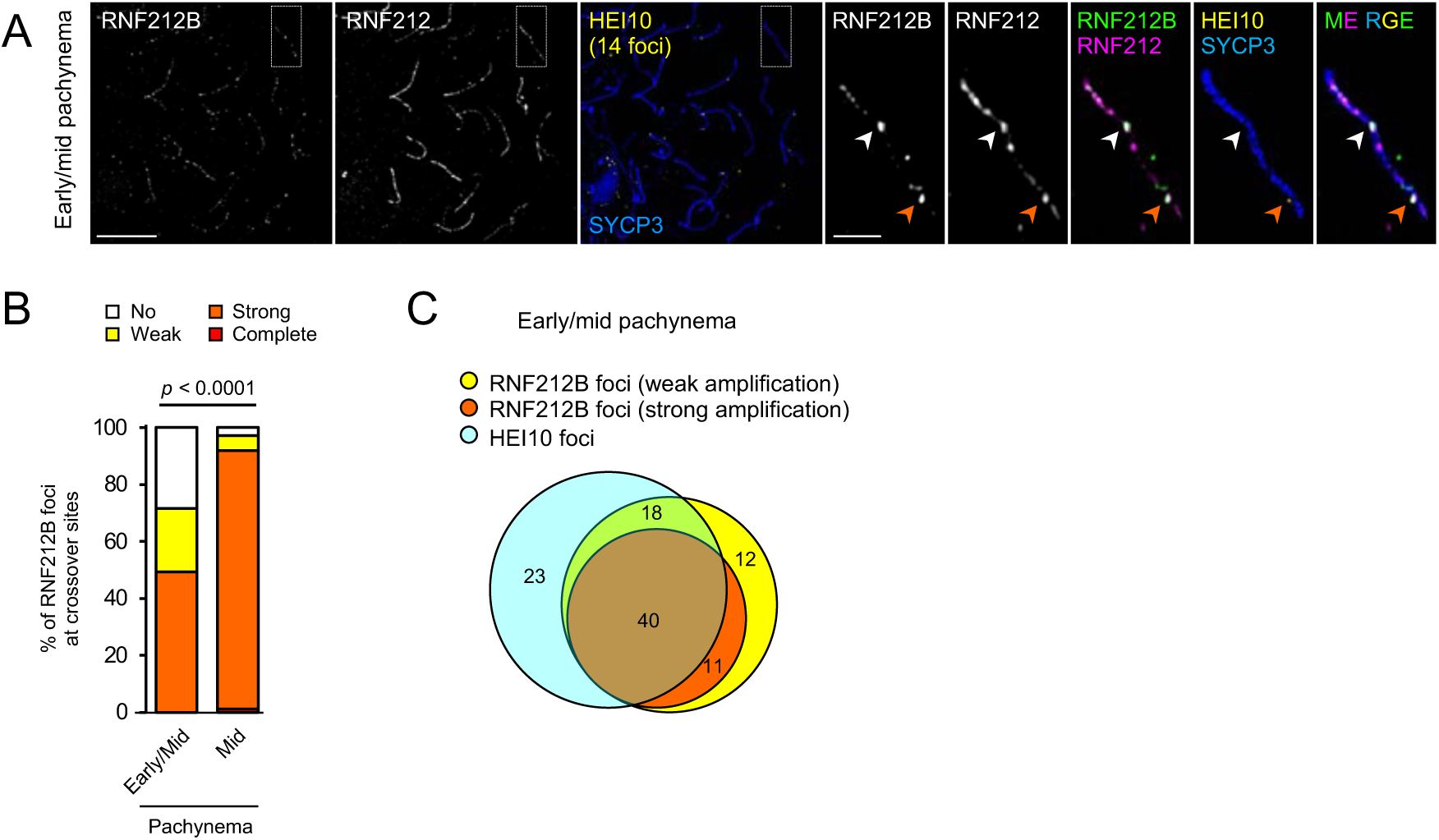
Timing of RNF212B focus amplification and appearance of HEI10 foci in spermatocytes, related to Figure 3. (A) Amplification of RNF212B foci relative to the emergence of HEI10 foci. Airyscan images of an early-to-mid pachytene spermatocyte nucleus immunostained for SYCP3, RNF212B, RNF212 and HEI10. This nucleus contains 14 HEI10 foci indicating that it is in transition to a full complement of designated crossovers sites (≥20 HEI10 foci). The white arrowhead indicates an amplified RNF212B focus without an associated HEI10 focus; and the orange arrowhead indicates an amplified RNF212B focus colocalized with a small, emerging HEI10 focus. These configurations indicate that growth of RNF212B foci can precede a detectable HEI10 focus. (B) Amplification of RNF212B at crossover sites. Crossover-specific foci of RNF212B (i.e. RNF212B foci that colocalized with HEI10 foci) in early-to-mid pachytene nuclei (containing 5-18 HEI10 foci), and mid-pachytene nuclei (≥19 HEI10 foci) were classified based on their degree of amplification by measuring their intensity relative to the brightest other (noncrossover) RNF212B focus along the same chromosome (complete means no other foci detected along the SC; strong, ≥2-fold brighter; weak, 1.5-2-fold brighter; or no amplification, <1.5-fold brighter). 81 crossover-specific RNF212B foci from 7 early-to-mid pachytene and 186 crossover-specific RNF212B foci from 8 mid-pachytene nuclei were analyzed by Airyscan imaging. In early-mid pachytene nuclei, the majority of HEI10 foci are associated with detectable amplification of RNF212B but 28% are not. (C) Amplification of RNF212B and emergence of HEI10 foci in early-to-mid pachynema. Numbers of HEI10 foci (sky blue), and RNF212B foci with weak (yellow) and strong amplification (orange) are shown. Criteria for RNF212B focus amplification are as in (B). 71.6% (58/81 foci) of sites marked by HEI10 were coincident with weak or strong amplification of RNF212B, while 78.4% (40/51 foci) of strongly amplified RNF212B foci colocalized with a HEI10 focus. Thus, with respect to the amplification of RNF212B and the emergence of HEI10 at prospective crossover sites, a sequential order of events is hard to discern, i.e. these events may be coincident, consistent with the dependence of RNF212B focus differentiation on HEI10. 7 early-to-mid pachytene nuclei with 5-18 HEI10 foci per nucleus were analyzed by Airyscan imaging.

**Figure S9.**
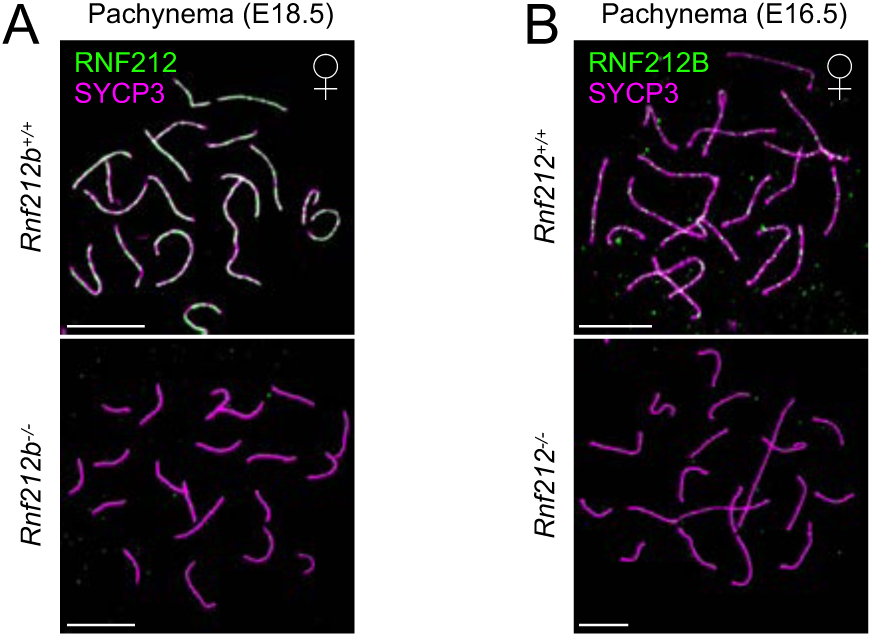
Interdependent chromosomal localization of RNF212B and RNF212 in oocytes, related to Figure 4. (A and B) RNF212 (A) and RNF212B (B) are interdependent for chromosomal localization. Pachytene-stage oocytes at embryonic day 18.5 or 16.5 (E18.5 or E16.5) from the indicated genotypes were immunostained for SYCP3 and RNF212 (A) or RNF212B (B). Scale bars, 10 μm.

**Figure S10.**
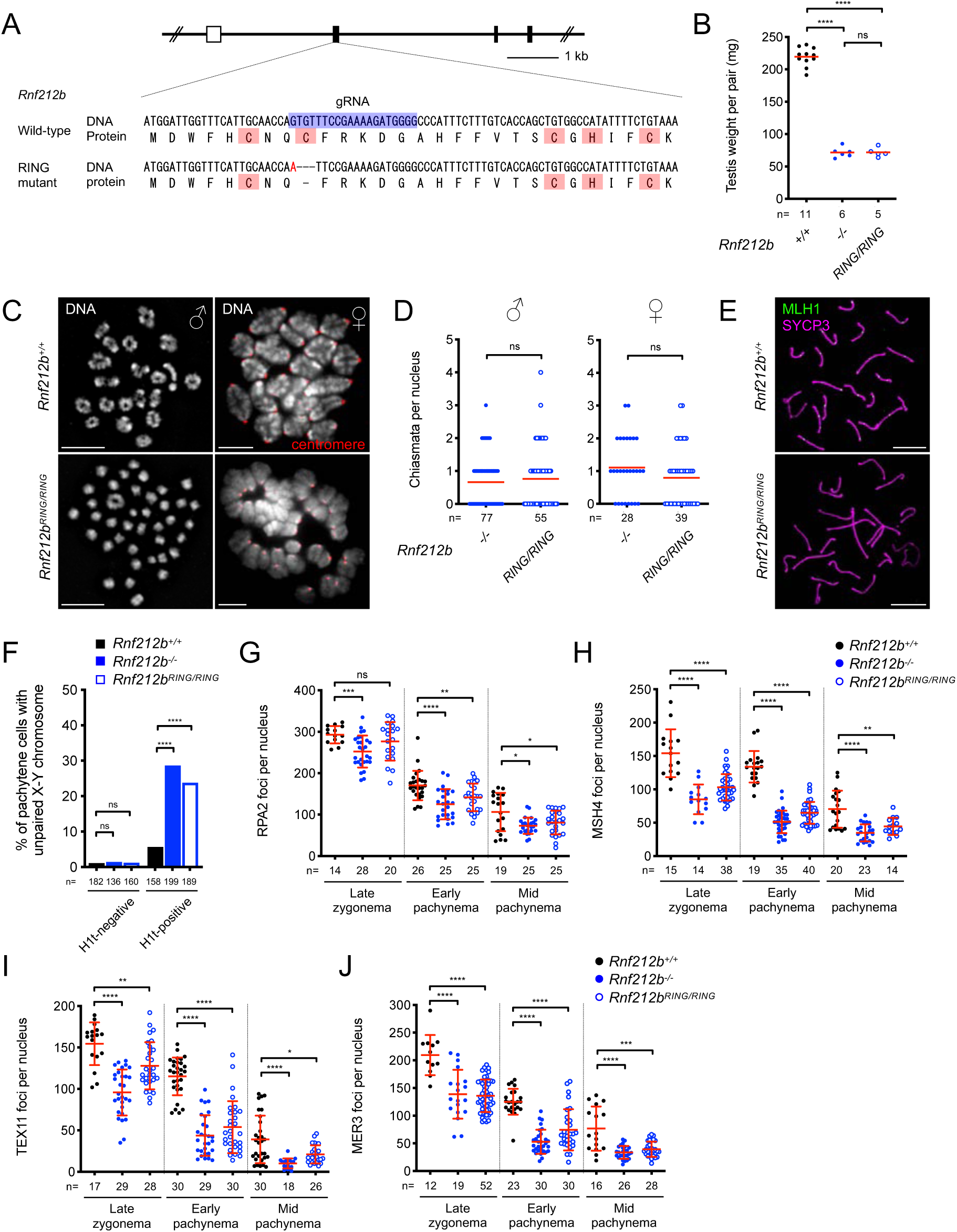
The RING finger domain is required for RNF212B function *in vivo*, related to Figure 4. (A) Scheme of the *Rnf212b* RING mutation. A three-nucleotides deletion in addition to a single-nucleotide substitution (shown in red) induced loss of the 9th cysteine residue with no change in other protein-coding amino acids. White and black boxes indicate 5’ UTR and coding exons, respectively. The guide RNA (gRNA) sequence is highlighted in blue. Cysteine and histidine residues of RING finger domain are highlighted in pink. (B) Reduced testis size in *Rnf212b^RING/RING^* mutants. Red bars indicate means. ns, not significant (*p* > 0.05); \*\*\*\**p* ≤ 0.0001 for two-tailed *t* tests. Numbers of mice analyzed are indicated below the X axis. (C) Diminished chiasma numbers in *Rnf212b^RING/RING^* meiocytes. Left, wild-type and *Rnf212b^RING/RING^* spermatocytes in diakinesis/metaphase-I stages stained with DAPI. Right, metaphase-I oocytes from ≥2 months old wild-type and *Rnf212b^RING/RING^* females were stained with DAPI and immunostained for centromeres. Scale bars, 10 μm. (D) Chiasma counts in *Rnf212b^-/-^* and *Rnf212b^RING/RING^* meiocytes. Red bars indicate means. ns, not significant (*p* > 0.05) for two-tailed Mann-Whitney tests. Total numbers of nuclei analyzed are indicated below the X axis. (E) Absence of MLH1 foci in *Rnf212b^RING/RING^* spermatocytes. Mid-late pachytene nuclei from wild-type and *Rnf212b^RING/RING^* spermatocytes immunostained for SYCP3 and MLH1. Scale bars, 10 μm. (F) High frequency of unpaired X-Y chromosomes in H1t-positive pachytene *Rnf212b^RING/RING^* cells. ns, not significant (*p* > 0.05); \*\*\*\**p* ≤ 0.0001 for Fisher’s exact tests. Total numbers of cells analyzed from 3 mice of each genotype are indicated below the X axis. (G-J) Focus counts of RPA2 (G), MSH4 (H), TEX11 (I), and MER3 (J) in wild-type, *Rnf212b^-/-^,* and *Rnf212b^RING/RING^* spermatocytes. Red bars indicate means ± SDs. ns, not significant (*p* > 0.05); \**p* ≤ 0.05; \*\**p* ≤ 0.01; \*\*\**p* ≤ 0.001; \*\*\*\**p* ≤ 0.0001 for two-tailed Mann-Whitney tests. Total numbers of nuclei analyzed are indicated below the X axes.

**Figure S11.**
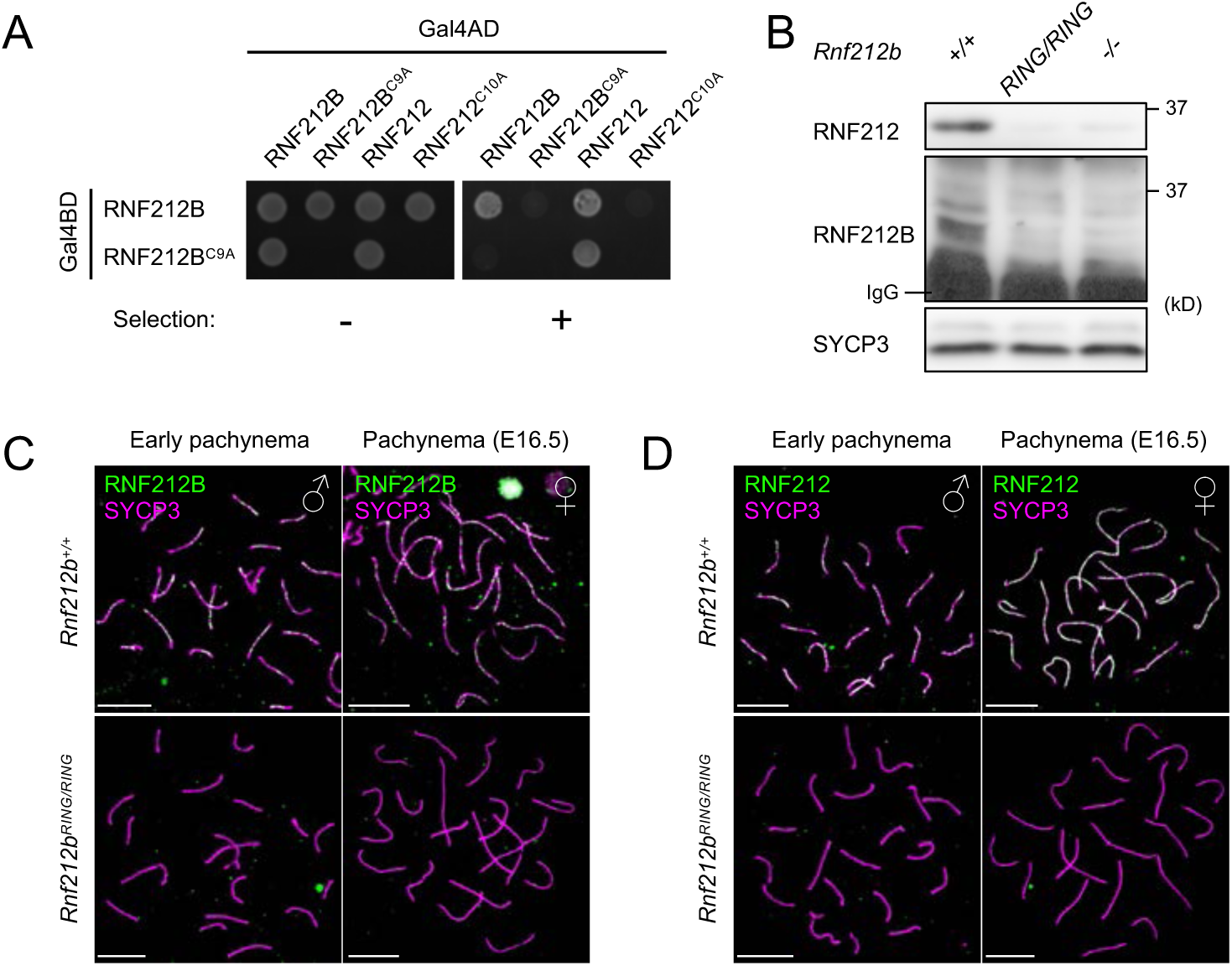
Role of the RNF212B RING-finger domain for self-interaction, chromosomal localization, and protein stability, related to Figure 4. (A) Yeast two-hybrid assay shows that a C9A RING-domain mutation in RNF212B diminishes its self-interaction but not its interaction with RNF212. (B) Diminished protein levels of RNF212B and RNF212 in *Rnf212b^RING/RING^* spermatocytes. Whole-testis extract from indicated genotypes were subjected to immunoblotting for RNF212 and IP-immunoblotting for RNF212B. SYCP3 is a loading control of meiotic cells. (C and D) Chromosomal localization of RNF212B (C) and RNF212 (D) is diminished in *Rnf212b^RING/RING^* mutant meiocytes. Early pachytene spermatocyte nuclei from wild-type and *Rnf212b^RING/RING^* testes (left); and pachytene-stage nuclei from wild-type and *Rnf212b^RING/RING^* oocytes at E16.5 (right) were immunostained for SYCP3 and RNF212B (C) or RNF212 (D). Scale bars, 10 μm.

**Figure S12.**
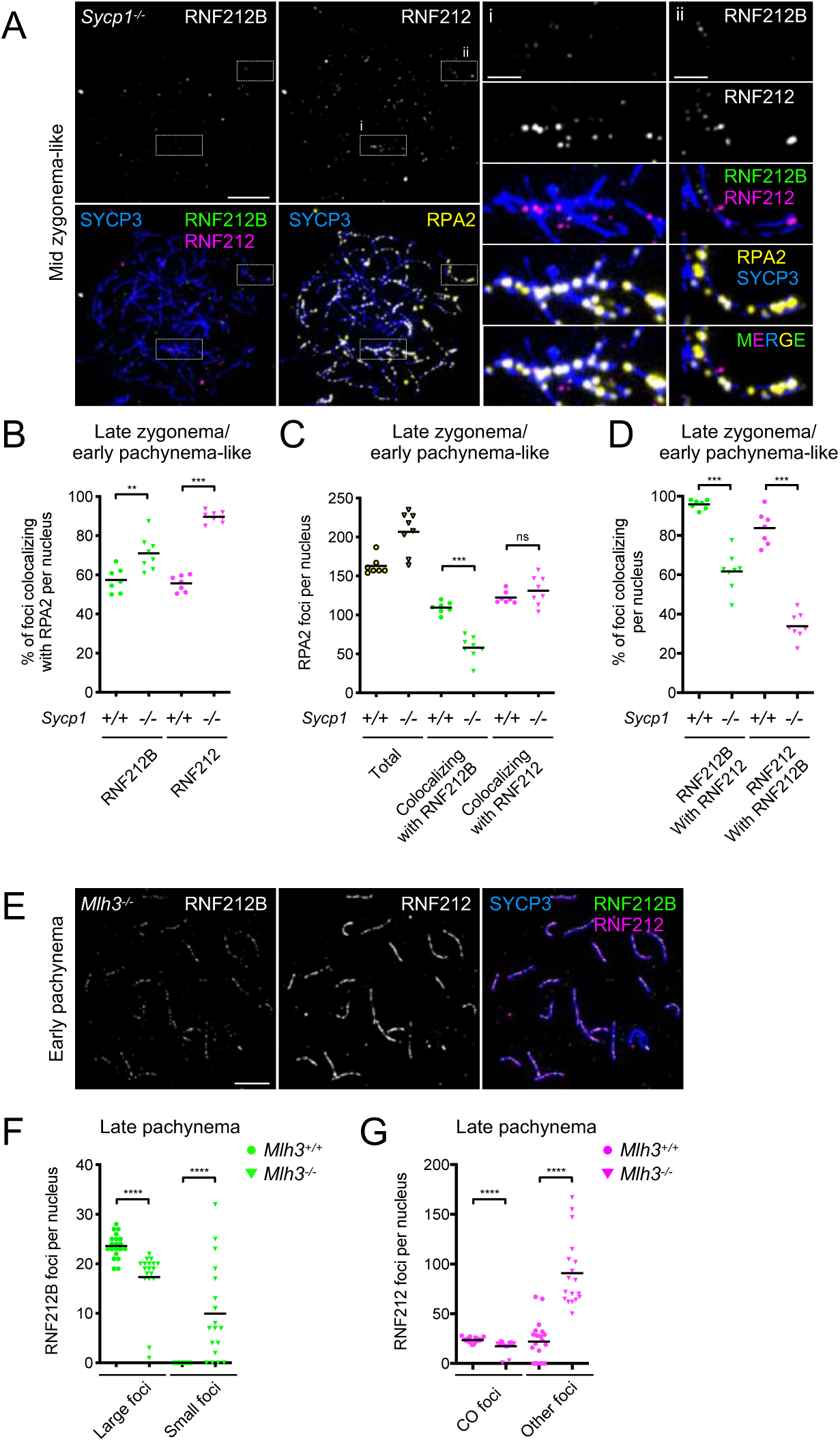
Chromosomal localization of RNF212B and RNF212 in mutants defective for synapsis and crossing over, related to Figure 5. (A) A mid zygotene-like *Sycp1^-/-^* spermatocyte nucleus immunostained for SYCP3, RNF212B, RNF212, and RPA2. General staining of RNF212B and RNF212 is absent but limited localization to small regions where homologous axes are closely aligned can be detected (magnified in the panels on the right). (B and C) Quantification of RPA2 foci colocalizing with RNF212B and RNF212. (B) Degree of RNF212B and RNF212 colocalization with RPA2. (C) Quantification of total and colocalizing RPA2 foci. Black bars indicate means. 7 early-pachytene *Sycp1^+/+^* nuclei and 8 late-zygotene/early-pachytene-like *Sycp1^-/-^* nuclei were analyzed. (D) Quantification of RNF212B-RNF212 colocalization. Black bars indicate means. 7 early pachytene *Sycp1^+/+^* nuclei and 8 late zygotene/early pachytene-like *Sycp1^-/-^* nuclei were analyzed. (E) Normal chromosomal localization of RNF212B and RNF212 in early pachytene *Mlh3^-/-^* mutant spermatocyte nuclei. An early pachytene nucleus immunostained for SYCP3, RNF212B and RNF212 is shown. (F) Numbers of RNF212B and RNF212 foci in wild-type and *Mlh3^-/-^* late-pachytene spermatocyte nuclei. Left, numbers of large and small RNF212B foci. Right, numbers of RNF212 foci associated with crossover sites (colocalizing with large RNF212B foci) and other sites. Black bars indicate means. 20 *Mlh3^+/+^* and 18 *Mlh3^-/-^* nuclei were analyzed. ns, not significant (*p* > 0.05); \*\**p* ≤ 0.01; \*\*\**p* ≤ 0.001; \*\*\*\**p* ≤ 0.0001 for two-tailed Mann-Whitney tests. Scale bars, 10 μm in images of full nuclei (A and E) and 2 μm in the magnified panels (A).

**Figure S13.**
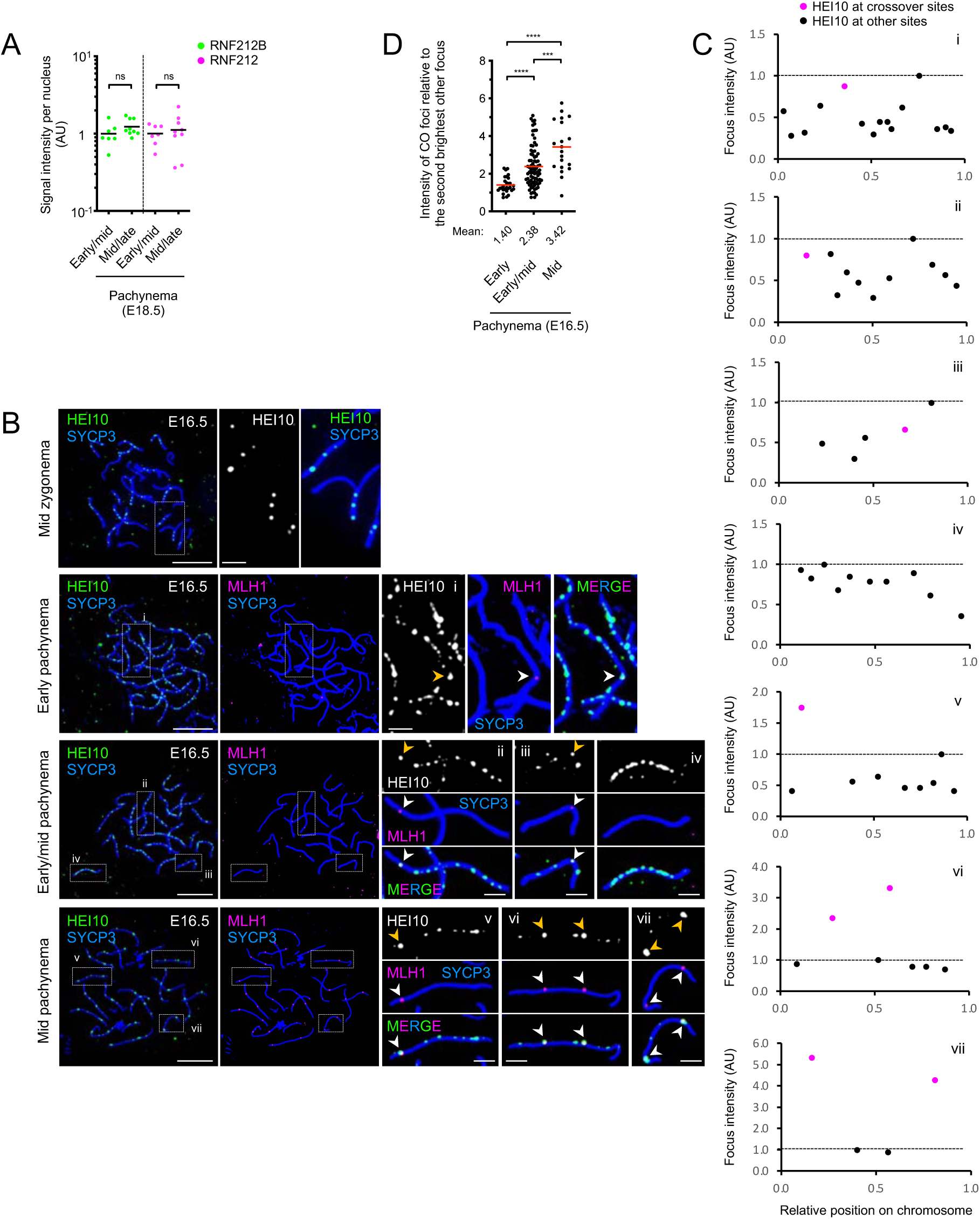
Localization of HEI10 and MLH1 in oocytes, related to Figures 6 and 7. (A) Total signal intensity of RNF212B and RNF212 per nucleus in early/mid and mid/late pachytene oocytes. Black bars indicate means. ns, not significant (*p* > 0.05) for two-tailed Mann-Whitney tests. Nuclei analyzed were the same as in Figure 6H. (B) HEI10 localization at successive prophase-I stages of fetal oocyte nuclei from embryonic day 16.5 (E16.5). Surface-spread prophase-I oocyte nuclei in mid zygonema, early pachynema, early/mid pachynema, and mid pachynema were immunostained for SYCP3, HEI10, and MLH1. Arrowheads indicate crossover sites marked by MLH1. The magnified images show representative chromosomal regions. Scale bars, 10 μm for images of full nuclei and 2 μm for magnified panels. (C) Representations of per chromosome analysis of HEI10 foci for the chromosomes highlighted in (B). Each HEI10 focus is represented by a dot. Magenta dots indicate HEI10 foci at crossover sites (colocalizing with MLH1 foci). The dashed line indicates the signal intensity of the brightest other focus (noncrossover focus) along the same chromosome. (D) Per chromosome analysis showing that HEI10 foci located at crossover sites (colocalizing with MLH1) increase in relative intensity as pachytene progresses. Mean intensities are indicated by red lines and values shown below the graph. ns, not significant (*p* > 0.05); \*\*\**p* ≤ 0.001 for two-tailed Mann-Whitney tests. 31, 91, and 105 crossover foci from 14 early pachytene, 9 early/mid pachytene, and 6 mid pachytene nuclei were analyzed, respectively.

## MATERIALS AND METHODS

### Mice

Mice were maintained and euthanized according to the guidelines of the Institutional Animal Care and Use Committees of the University of California Davis. All mice were congenic with the C57BL/6J background except for *Sycp1^-/-^* mutant mice. The *Rnf212^-/-^*, *Hei10 ^mei^*^4^*^/mei4^*, *Spo11^-/-^*, *Sycp1^-/-^* and *Mlh3^-/-^* mutant lines were previously described ^15, 25, 49, 51, 52^. Mature adult mice (2-6 months old) were used for experiments unless otherwise noted.

*Rnf212b* mutant alleles were generated by The Cornell Stem Cell and Transgenic Core Facility. 124 bp of DNA template for sgRNA, comprising the T7 polymerase binding site and sgRNA target sequence (5’-CCCCATCTTTTCGGAAACAC-3’) followed by the remaining sgRNA sequence, were amplified by PCR as previously described ^63^ and purified using a GeneJET PCR Purification Kit (Thermo Scientific, K0702). 600 ng of purified DNA template was *in vitro* transcribed for 4 h at 37°C using MEGAshortscript T7 Transcription Kit (Invitrogen, AM1354). sgRNA was purified using MEGAclear Transcription Clean-Up Kit (Invitrogen, AM1908), diluted to 1.6 μg/μl and stored at -80°C prior to embryo injection. 12.5 ng/μl of purified sgRNA and either 12.5 or 25 ng/μl of Cas9 mRNA were microinjected into pronuclei of 2-cell stage of C57BL/6J homozygous embryos. Embryos were transferred to pseudo-pregnant female recipients 2 days after pronuclear injection at the 4-cell stage. Genomic DNA of founder mice was isolated from toe clips, PCR-amplified, cloned into the pCR4Blunt-TOPO vector (Invitrogen, 450031), and Sanger-sequenced to identify mutations. Founder mice were backcrossed to C57BL/6J mice for ≥4 generations and heterozygous mice were bred to homozygosity. Genotyping was performed by PCR on genomic DNA isolated from mouse tails. Primers used for cloning and genotyping are listed in **Table S2**.

### Total RNA extraction, RT-PCR, and 5’ RACE

Tissues were dissected from adult male mice, washed in phosphate-buffered saline (PBS), frozen in liquid nitrogen and stored at -80°C prior to RNA extraction. Total RNA was extracted using TRIzol according to manufacturer’s instructions (Invitrogen). 1 μg of total RNA was reverse-transcribed using the SuperScript IV First-Strand Synthesis System (Invitrogen, 18091050) and synthesized cDNA was PCR-amplified. For 5’ RACE, 4 μg of total RNA from testes was reverse-transcribed using 5’ RACE System for Rapid Amplification of cDNA Ends (Invitrogen, 18374058) and synthesized cDNA was PCR-amplified, cloned into the pCR4-TOPO TA vector (Invitrogen, K457501) and Sanger-sequenced to identify translation initiation site. Primers used are listed in **Table S2**.

### Yeast two-hybrid assays

Full-length mouse *Rnf212b* and *Rnf212* cDNAs were PCR amplified from mouse testis cDNA, prepared as above, and cloned into pGADT7 and pGBKT7 vectors (Clontech) using a Gibson Assembly Cloning Kit (NEB, E2611S). Full-length mouse *Hei10* coding sequence was synthesized by Twist Bioscience and cloned into pGADT7 and pGBKT7. Prey and bait vectors were co-transformed into the Y2HGold strain. Transformants were suspended in 20 μl sterile water, and 2 μl spotted onto fresh plates with and without selection: SD/-Trp/-Leu plate, SD/-Trp/- Leu containing 200 ng/ml of Aureobasidin A (Clontech, 630466) to select for the *AUR1-C* reporter gene, or SD/-Trp/-Leu /-Ade/-His to select for *ADE2* and *HIS3* reporters. After incubation of the plates at 30°C for 3 days, single colonies were inoculated into SD/-Trp/-Leu media and grown overnight. Colonies were resuspended at OD_600_ = 1, four 10-fold serial dilutions were prepared, and 10 μl drops were spotted onto selection plates and incubated at 30°C for 3 to 5 days. Primers used for cloning and point mutagenesis are listed in **Table S2**.

### Antibody production

Polyclonal antibodies against mouse RNF212B were raised in Guinea pigs. Codon-optimized of full-length mouse RNF212B was cloned into pET-28b (+) (Addgene) with a C-terminal 6xHis tag. *E. coli* Arctic Express (DE3) cells (Agilent Technologies Inc.) transformed with the mouse RNF212B-6xHis expression vector were grown in 2L of LB at 30°C to an OD_600_ of 0.8, and protein expression was induced with 0.5 mM IPTG followed by incubation at 11°C in media supplemented with 0.1 mM ZnCl_2_. ∼5 g of cells were pelleted by centrifugation, suspended in 60 ml of denaturing lysis buffer A (6M Guanidine, 25 mM sodium-phosphate pH 7.4, 500 mM NaCl, 0.1 mM ZnCl_2_, 1 mM β-mercaptoethanol, 20 mM imidazole, 10% glycerol) and sonicated. Samples were centrifuged at 35,000 rpm in a Ti-45 rotor (Beckman) for 45 min and soluble extract was applied to a 5 ml HisTrap FF column (GE healthcare Inc.) using a GE AKTA Avant 25 FPLC system. After equilibrating with 5 column volumes (CV) of lysis buffer A, followed by washes with 10 CV of wash buffer B (6M urea, 25 mM sodium-phosphate pH 7.4, 500 mM NaCl, 1 mM β-mercaptoethanol, 10% glycerol) supplemented with 60 mM imidazole, bound proteins were eluted with a linear gradient of imidazole (60-600 mM) in buffer B. Peak fractions were pooled and dialyzed extensively in decreasing urea concentrations (6M to 1M urea in 25 mM sodium-phosphate pH 7.4, 500 mM NaCl, 1 mM β-mercaptoethanol, 10% glycerol). Protein concentrations were determined by Bradford and spectrophotometric (A_280_) methods. Two Guinea pigs were immunized with purified RNF212B protein by Antibodies Incorporated (Davis, CA). The IgG fraction was purified from the resultant sera, using the Montage antibody purification kit according to manufacturer’s instructions (Millipore Sigma), and dialyzed against PBS containing 10% glycerol and aliquots stored at -80°C.

### Histology

Testes from adult male mice were dissected, punctured with a 27-guage needle, and fixed in 10% buffered formalin overnight at room temperature. Ovaries from 18 days postpartum (18 dpp) female mice were dissected and fixed in 10% buffered formalin overnight at room temperature. Fixed testes and ovaries were washed in PBS twice and stored in 70% ethanol at 4°C prior to embedding. Tissues embedded in paraffin were sectioned (5 μm) onto glass slides (Fisher Scientific, 12-550-15), deparaffinized, rehydrated, and incubated with antigen retrieval buffer (10 mM Sodium Citrate, 0.05% Tween-20) for 50 min at 100°C. Slides were then stained with hematoxylin and eosin (testes), or immunostained with anti-p63 antibody, as described below, and counterstained with hematoxylin (ovaries), and mounted with Permount (Fisher Scientific, SP15-100).

### Surface spreads of spermatocyte chromosomes

Surface-spread chromosomes of spermatocytes were prepared as described previously ^64^with slight modification. Testes were dissected, the tunica albuginea removed, and adherent extratubular tissues removed by rinsing and dissociating seminiferous tubules in PBS using a pair of 25-gauge (adult) or 27-guage (juvenile) needles in a 35 mm petri dish (Corning, 430165). Dissociated seminiferous tubules were incubated in hypotonic extraction buffer (30 mM Tris-HCl pH 8.0, 50 mM sucrose, 17 mM trisodium citrate dihydrate, 5 mM EDTA, 0.5 mM dithiothreitol (DTT) and 0.5 mM phenylmethylsulphonyl fluoride (PMSF), pH 8.2-8.3) for 15-45 min at room temperature. After quickly rinsing seminiferous tubules in 100 mM sucrose, a small amount of tubules were placed in 40 μl of 100 mM sucrose on a glass depression slide. Tubules were torn to pieces using two fine forceps, and large pieces of tubular remnant were removed. The volume was increased to 40-60 μl by adding 100 mM sucrose and a cell suspension was made by pipetting. A clean glass slide (Fisher Scientific, 12-544-7) was dipped into freshly made PFA solution (1% paraformaldehyde adjusted to pH 9.2 using 1.25 M sodium borate, 0.15% Triton X-100) in a 50 ml Falcon tube, and excess solution was drained onto a paper towel. 20 μl of cell suspension was placed at the upper right corner of the slide and slowly dispersed in horizontal and vertical directions to homogeneously cover the slide. Hot tap water was added to a humid chamber and slides were slowly dried in the closed chamber overnight at room temperature. Slides were further dried for 3 hr with lid ajar, and then for 1-2 hr with the lid removed. Slides were washed once for 5 min in deionized water and twice for 5 min in 0.4% Photo-Flo 200 solution (Kodak, 1464510) in a coplin jar before air-drying at room temperature. Slides were either directly processed for immunostaining, as described below, or stored wrapped in aluminum foil at -80°C prior to immunostaining. For SIM imaging, 15 μl of cell suspension was placed at the upper right corner of a coverslip (Fisher Scientific, 12-544-E) coated with 90 μl of 1% PFA, 0.15% Triton X-100 solution and slowly dispersed as above.

### Surface spreads of oocyte chromosomes

Surface-spread oocyte chromosomes were prepared as described for spermatocytes with modifications. Ovaries from fetal females were dissected into PBS and kept on ice prior to incubation in hypotonic extraction buffer. A pair of ovaries were incubated in hypotonic extraction buffer for 10 min at room temperature, quickly rinsed in 100 mM sucrose, and placed in 50 μl of 100 mM sucrose on a glass depression slide. Ovaries were torn to pieces using a pair of 25-guage needles, and large pieces of ovarian remnant were removed. The volume was increased to 40 μl by adding 100 mM sucrose and a cell suspension was made by pipetting. One half of a clean glass slide was coated with 50 μl of freshly made 1% PFA 0.15% Triton X-100, and 10 μl of cells suspension was placed at the upper right corner of the slide slowly dispersed in horizontal and vertical directions to homogeneously cover the half slide. Slides were then dried, washed, and stored as described for spermatocyte chromosomes spreads.

### Chromosome spreads of diakinesis/metaphase-I spermatocytes

Testes from adult male mice were dissected, tunica albuginea removed, and seminiferous tubules placed in 2 ml of hypotonic solution (1% trisodium citrate) in a 35 mm petri dish. Seminiferous tubules were torn to pieces using two fine forceps and, after adding 1 ml of hypotonic solution, 3 ml of tubule suspension was transferred to a 15 ml Falcon tube using a plastic transfer pipette (Phenix, PP-137030). Tubule fragments were allowed to settle out for 3 min and the supernatant containing suspended cells was transferred to another 15 ml Falcon tube. Remnant tubule fragments in the petri dish were suspended in 2 ml of hypotonic solution and transferred to the first 15 ml Falcon tube containing tubule fragments. The mixture of tubule fragments was resuspended using a transfer pipette, tubule fragments were settled out, and the supernatant was transferred to the 15 ml Falcon tube containing the first supernatant. The mixture of supernatants was filtered through 70 μm and 40 μm Cell Strainers (Corning 352350; Celltreat, 229481) and suspended cells were pelleted by centrifugation at 900 rpm for 10 min at room temperature. Cells were fixed by adding 3 ml of freshly made fixative solution 1 (75% methanol, 25% acetic acid with 0.375% chloroform) drop-by-drop while gentle vortexing. Cells pelleted by centrifugation at 900 rpm for 10 min at room temperature, resuspended in 3 ml of freshly made fixative solution 2 (75% methanol, 25% acetic acid, chilled to -20°C), pelleted again, and resuspended in 0.5 ml of fixative solution 2. Fixed cell suspension was dropped onto a clean glass slide (Fisher Scientific, 12-544-7) from a height of 2-3 feet height using a glass Pasteur pipette in a room with >40% humidity. Slides were air-dried for 10 min at room temperature and either stained immediately with ProLong Gold or Diamond Antifade Mountant (Thermo Fisher Scientific, P36930 or P36970) containing 1 μg/ml DAPI, or stored at 4°C prior to staining.

### Chromosome spreads of metaphase-I oocytes

Ovaries from adult female mice without prior hormonal stimulation were dissected, follicles were punctured in pre-warmed (37°C) M2 media (Sigma-Aldrich, M7167) using a 25-gauge needle, and germinal-vesicle stage oocytes with integral cumulus cell layers were collected. Surrounding cumulus cells were mechanically removed by pipetting and oocytes were cultured in M2 media for 7 hr at 37°C. Metaphase-I oocytes were transferred into Tyrode’s acidic solution (Sigma-Aldrich, T1788) to remove the zona pellucida (ZP), and ZP-free oocytes were transferred and maintained in M2 media prior to spreading. 5-10 μl of 1% PFA solution pH 9.2 containing 0.15% Triton X-100 and 3 mM DTT was placed in each well of a 12-well glass slide (Electron Microscopy Sciences, 63425-05) and one ZP-free oocyte was transferred to each well. Slides were air-dried overnight at room temperature, and either directly processed for immunostaining or stored at 4°C for up to several days prior to immunostaining.

### Immunostaining

Slides were rehydrated with Tris-buffered saline (TBS) pH 8.0 containing 0.05% of Triton X-100 (TBST) for 3 min, blocked twice with blocking buffer (1% normal goat or donkey serum, 3% bovine serum albumin (BSA), 1x TBS pH 8.0, 0.05% Triton X-100, 0.05% sodium azide) for 15 min at room temperature and incubated with primary antibodies in antibody dilution buffer (10% normal goat or donkey serum, 3% BSA, 1x TBS pH 8.0, 0.05% Triton X-100, 0.05% sodium azide) in a humid chamber overnight at room temperature. Slides were briefly rinsed with TBST, washed twice with TBST for 5 min, blocked twice with blocking buffer for 15 min at room temperature and then incubated with secondary antibodies in antibody dilution buffer in a humid chamber for 1 hr at 37°C. Slides were rinsed with TBST, washed three times with TBST for 5 min, washed once with Milli-Q water for 2 min, and air-dried prior to mounting with ProLong Gold or Diamond Antifade Mountant. For SIM imaging, cover slips were blocked three times for 15 min prior to primary antibody incubation.

For metaphase-I oocytes, slides were briefly rehydrated with TBST, washed three times with TBST for 5 min, blocked twice with blocking buffer for 15 min at room temperature, and incubated with primary antibodies in antibody dilution buffer in a humid chamber overnight at room temperature. Slides were rinsed with TBST, briefly washed twice with TBST, blocked twice with blocking buffer for 15 min at room temperature, and incubated with secondary antibodies in antibody dilution buffer in a humid chamber for 1 hr at room temperature. Slides were rinsed with TBST, briefly washed with TBST, and then washed three times with TBST for 5 min. Slides were then incubated with TBST containing 5 μg/ml DAPI for 10 min at room temperature and mounted with 50% ProLong Gold or Diamond Antifade Mountant in TBST. All primary and secondary antibodies used are listed in **Table S3**.

### Protein blot analysis

Testes were dissected from 16-18 dpp juvenile male mice and frozen in liquid nitrogen. Five pairs of testes were homogenized in 1 mL ice-cold RIPA buffer (50 mM Tris-HCl pH 7.5, 150 mM NaCl, 1 mM EDTA, 1% NP-40, 0.5% sodium deoxycholate, 0.1% sodium dodecyl sulfate, SDS) supplemented with protease and isopeptidase inhibitors (1x Complete protease inhibitor EDTA-free, Roche, 04693159001), 1 mM PMSF, 10 mM N-ethylmaleimide, NEM), incubated for 15 min on ice, sonicated (20% duty cycle, output 2 in burts for 2.5 min), and then incubated for 15 min on ice. Following centrifugation at 14,000 rpm for 15 min at 4°C, supernatants were collected as whole-testis extracts. Protein concentrations determined by Bradford assay were normalized, and samples were subjected to electrophoresis and immunoblotting. Membranes were blocked with TBST plus 2.5% non-fat milk for 1 hr at room temperature and incubated with primary antibodies overnight at 4°C. Membranes were washed three times with TBST for 10 min each, then incubated with HRP-conjugated secondary antibodies in TBST for 1 hr at room temperature. After four washes with TBST for 10 min each, HRP signal was developed using the SuperSignal West Pico Chemiluminescent Substrate (Thermo Scientific, PI-34080) and detected using an Amersham Imager 600 (GE Healthcare).

All primary and HRP-conjugated secondary antibodies used are listed in **Table S3**.

### Image acquisition

Images of surface-spread prophase chromosomes and metaphase-I chromosome spreads were acquired using a Zeiss AxioPlan II microscope with a 63 x Plan-Apochromat 1.4 NA objective and EXFO X-Cite metal halide light source, captured with a Hamamatsu ORCA-ER CCD camera and processed using Volocity (Perkin Elmer) and Photoshop (Adobe) software. SIM images were acquired using a Nikon N-SIM super-resolution microscope and processed using NIS-Elements 2 image processing software. Airyscan images were acquired using a Zeiss LSM800 with Airyscan microscope with a 60 x 1.4 NA objective and processed using ZEN imaging software (Carl Zeiss). Images of testis sections were acquired using a Zeiss Axio Imager M2 microscope with a 20 x Plan-Apochromat 0.8 NA objective, captured with a Hamamatsu ORCA-Flash 4.0 V3 sCMOS camera and processed using ZEN imaging software (Carl Zeiss). Images of ovary sections were acquired using a ScanScope digital scanner (Asperio) and processed using ImageScope software.

### Image analysis

Comparisons were made between animals that were either littermates or matched by age. Numbers and colocalization of foci were determined manually. *Rnf212b^-/-^*, *Rnf212^-/-^* and *Rnf212b^RING/RING^* mutant analysis was blinded with respect to the genotype of the animals. Results from more ≥2 independent experiments/animals were pooled for quantification of number of foci and chiasmata. *Spo11^-/-^*, *Sycp1^-/-^*, *Mlh3^-/-^* and *Hei10^mei4/mei4^* mutant analysis was not blinded to genotype because phenotypes are overt. For focus intensity analyses, RNF212B and RNF212 foci along homolog axes were automatically defined by thresholding intensity on SYCP3 staining and manually inspecting images to separate any adjacent focus pairs defined as one focus. For *Spo11^-/-^* mutant analysis, synapsed regions marked by SYCP1 were manually cropped and intensities of RNF212B and RNF212 were measured. Two regions of interest (ROI) were drawn inside nuclei but adjacent to homolog axes and their average intensity was subtracted as background for each nucleus. Results from one experiment/animal among ≥2 independent experiments/animals were shown for quantification of intensities. Prophase-I stages were defined by SYCP3 staining (spermatocytes) and both SYCP3 staining and fetal age (oocytes) using standard criteria. Leptonema was defined by short SYCP3 stretches without evidence of synapsis determined by thickening of SYCP3 staining. Zygonema was defined by longer stretches of SYCP3 with various degree of synapsis: early, mid- and late-zygonema were defined by having <25%, 25-75% and >75% of synapsis, respectively. Pachynema was defined by full synapsis of all autosomes. For spermatocytes, early pachynema was defined by extensive synapsis between the X-Y chromosomes, mid pachynema was defined by limited/end-to-end X-Y synapsis, and late pachynema was defined by decondensed/elongated X-Y chromosomes with figure-of eight configuration and thickening of SYCP3 staining at telomeres of autosomes. Diplonema in spermatocytes was defined by internal desynapsis with various degrees of residual synapsis: early, mid and late diplonema were defined by having >75%, 75-25% and<25% of synapsis, respectively. For oocytes, nuclei with unsynapsed/desynapsed chromosomes at embryonic day 15.5 (E15.5) and at 0.5 day postpartum (0.5 dpp) were defined as zygonema and diplonema, respectively. Dictyate stage was defined by short, ragged SYCP3 stretches without evidence of synapsis at 0.5 dpp.

For ovary sections, numbers of oocytes were manually counted for every fifth section and counts were multiplied by five to calculate the total number of oocytes per pair of ovaries, as described previously ^65^.

### Statistical analysis

Statistical analyses were performed using Graphpad Prism software v.8-9 and R version 3.5.2. Statistical parameters and tests, sample sizes, means, error bars, and standard deviations (SDs) are described in the figures and/or corresponding figure legends. Sample sizes were not predetermined using any statistical tests.

## Acknowledgements

We thank M.A. Handel, P. Cohen, C. Höög, J. Ward, and R.J. Pezza for antibodies; the UC Davis MCB Light Microscopy Imaging Facility and M.R. Paddy for help with SIM imaging; the Cornell Stem Cell and Transgenic Core Facility, supported in part by Empire State Stem Cell Fund Contract Number C024174, J. Schimenti and R.J. Munroe for generating *Rnf212b* mutant mice; and members of the Hunter lab for helpful discussions. M.I. was supported by a Japan Society for the Promotion of Science postdoctoral fellowship for research abroad. S.S. was supported by A.P. Giannini postdoctoral fellowship. B.N. was supported by the Howard Hughes Medical Institute EXROP program. N.H. is an Investigator with the Howard Hughes Medical Institute.

## Author contributions

M.I. and N.H. conceived the study and designed the experiments. M.I., S.L., L.H., K.L., N.L., R.M.H., R.P., C.H., and I.H. performed the experiments and analyzed the data. Y.Y. performed oocyte culture experiments and analyzed the data. S.S. and B.N. performed histological experiments and analyzed the data. D.S.K. generated anti-RNF212B antibodies. M.I. and N.H. wrote the manuscripts with input from Y.Y., S.L., S.S., and D.S.K. All authors edited the manuscript.

