## Supplemental Tables for "Distinct and interdependent functions of three RING proteins regulate recombination during mammalian meiosis" (from another version of this preprint: version 2, 2024-09-15)

**Table S1. Mutations and phenotypes of *Rnf212b* mutant lines generated in this study.**

Founder mice with mono-allelic mutation

| ID | Mutation | Homozygous mutant male |
|---|---|---|
| 1E | 1 nt insertion after 26 nt from first ATG / a premature stop codon after 27 a.a. | Infertile, small testes |
| 2Q | 1 nt insertion after 26 nt from first ATG / a premature stop codon after 27 a.a | Infertile, small testes |
| 3K | 1 nt substitution after 23 nt from first ATG / no change in coding a.a.; | Infertile, small testes |
| | 3 nt deletion after 24 nt from first ATG / C9- mutation | |
| 4A | 10 nt deletion after 23 nt from first ATG / a premature stop codon after 27 a.a. | Infertile, small testes |
| 5C | 1 nt insertion after 27 nt from first ATG / a premature stop codon after 27 a.a. | Infertile, small testes |
| 6U | 8 nt insertion after 26 nt from first ATG / a premature stop codon after 8 a.a. | Infertile, small testes |
| 7W | 1 nt insertion after 26 nt from first ATG / a premature stop codon after 8 a.a. | Infertile, small testes |

Founder mice with bi-allelic or chimeric mutation

| ID | Allele | Mutation | Founder mouse |
|---|---|---|---|
| 8M | 1 | 2 nt deletion after 25 nt from first ATG / a premature stop codon after 26 a.a. | Infertile, small testes |
| | 2 | 2 nt deletion followed by 1 nt substitution after 26 nt from first ATG / a premature stop codon after 26 a.a. | |
| 9P | 1 | 1-5-1 nt deletion after 16 nt from first ATG / a premature stop codon after 28 a.a. | Infertile, small testes |
| | 2 | 2 nt insertion after 27 nt from first ATG / a premature stop codon after 31 a.a. | |
| | 3 | 3 nt insertion followed by 1 nt substitution after 25 nt from first ATG | |
| 10G | 1 | 1 nt insertion after 26 nt from first ATG / a premature stop codon after 27 a.a. | Infertile (female) |
| | 2 | 354 nt insertion after 25 nt from first ATG | |

| | | | |
|---|---|---|---|
| 11I | 1 | 9 nt deletion after 22 nt from first ATG / Q8-, C9-, F10- mutation | Infertile (female) |
| | 2 | 12 nt deletion after 27 nt from first ATG / F10-, R11-, K12-, D13- mutation;<br>1 nt substitution after 61 nt from first ATG / S21N mutation | |
| 12Y | 1 | 2 nt deletion after 25 nt from first ATG / a premature stop codon after 26 a.a. | Infertile (female) |
| | 2 | 4 nt deletion after 22 nt from first ATG / a premature stop codon after 29 a.a. | |

**Table S2. Primers used in this study**

| **Primer** | **Sequence (5'->3')** |
|---|---|
| Cloning of *Rnf212b* gene to identify mutation, forward | GTTCTCTGGCATATAGTGAAG |
| Cloning of *Rnf212b* gene to identify mutation, reverse | CCCGACACAACTCAAAAGAC |
| Genotyping of *Rnf212b* wild-type allele, forward | TGGTTTCATTGCAACCAGTGT |
| Genotyping of *Rnf212b* null mutant allele, forward | TGGTTTCATTGCAACCAGTGG |
| Genotyping of *Rnf212b* RING mutant allele, forward | GGTTTCATTGCAACCAATTCC |
| Genotyping of *Rnf212b*, common reverse | CCCGACACAACTCAAAAGAC |
| RT-PCR for *Rnf212b* 5'UTR variant a, forward | CTGCTTCTCAAATGTCTATAAAG |
| RT-PCR for *Rnf212b* 5'UTR variant b, forward | CTGCTTCTCAAATGTAACAAGG |
| RT-PCR for *Rnf212b* 5'UTR variants a/b/c, forward | CTGCCAAGCTGAATGGATTG |
| RT-PCR for *Rnf212b* 5'UTR variants, common reverse | GAGAAGGGGTTTCAGTGTAAG |
| RT-PCR for *Rnf212b* isoforms a, b, c, forward | CTTTGGAAAAATGTGCTGTTTG |
| RT-PCR for *Rnf212b* isoform d, forward | CACATCTCTCAGTAGGTGTG |

| | |
|---|---|
| RT-PCR for *Rnf212b* isoforms a, d, reverse | GAGAAGGGGTTTCAGTGTAAG |
| RT-PCR for *Rnf212b* isoform b, reverse | GAGTCTTTCAGGCAGTGTAAG |
| RT-PCR for *Rnf212b* isoform c, reverse | TGGAGAAGGGGTTTCTGTTC |
| Cloning of *Rnf212b* cDNA into pGADT7, forward | ATGGCCATGGAGGCCAGTGAATTCATGGATTGGTTTCATTGCAACCAG |
| Cloning of *Rnf212b* cDNA into pGADT7, reverse | TGCAGCTCGAGCTCGATGGATCCCTATCTGGAAATGTTCCATCCGTCTTT |
| Cloning of *Rnf212b* cDNA into pGBKT7, forward | CATATGGCCATGGAGGCCGAATTCATGGATTGGTTTCATTGCAACCAG |
| Cloning of *Rnf212b* cDNA into pGBKT7, reverse | GGCCGCTGCAGGTCGACGGATCCCTATCTGGAAATGTTCCATCCGTCTTT |
| Cloning of *Rnf212* cDNA into pGADT7, forward | ATGGCCATGGAGGCCAGTGAATTCATGGCCAGCTGGGTGTTCTGTAAT |
| Cloning of *Rnf212* cDNA into pGADT7, reverse | TGCAGCTCGAGCTCGATGGATCCTCATCGATGAACGCATGCATGCCAGGG |
| Cloning of *Rnf212* cDNA into pGBKT7, forward | CATATGGCCATGGAGGCCGAATTCATGGCCAGCTGGGTGTTCTGTAAT |
| Cloning of *Rnf212* cDNA into pGBKT7, reverse | GGCCGCTGCAGGTCGACGGATCCTCATCGATGAACGCATGCATGCCAGGG |
| Point mutagenesis for $Rnf212b^{C9A}$ | TTGGTTTCATTGCAACCAGGCTTTCCGAAAAGATGGGGCC |
| Point mutagenesis for $Rnf212^{C10A}$ | GGTGTTCTGTAATCGCGCTTTCCAGTCGCCGCAC |

**Table S3. Antibodies used in this study**

| Antibody | Source | Identifier | Application | Dilution |
|---|---|---|---|---|
| Guinea pig anti-RNF212B | this study | N/A | IF | 1:40 (SIM, 1:20) |
| | | | IB | 1:500 |
| Goat anti-RNF212 | Bondarieva et al., 2020 | N/A | IF | 1:50 (SIM, 1:25) |
| Guinea pig anti-RNF212 | this study | N/A | IB | 1:1,000 |

| | | | | |
|---|---|---|---|---|
| Mouse anti-SYCP3 | Santa Cruz Biotechnology | sc-74569 | IF | 1:200 (SIM, 1:50) |
| Goat anti-SYCP3 | Santa Cruz Biotechnology | sc-20845 | IF | 1:200 (SIM, 1:100) |
| | | | IB | 1:1,000 |
| Rabbit anti-SYCP3 | Santa Cruz Biotechnology | sc-33195 | IF | 1:200 |
| Rabbit anti-SYCP1 | Abcam | ab15090 | IF | 1:200 |
| Mouse anti-MLH1 | BD Pharmingen | 550838 | IF (spermatocyte) | 1:50 |
| Mouse anti-MLH1 | Cell Signaling Technology | 3515S | IF (oocyte) | 1:30 |
| Rabbit anti-HEI10 (serum) | Qiao et al., 2014 | N/A | IF | 1:500 |
| Mouse anti-CDK2 | Santa Cruz Biotechnology | sc-6248 | IF | 1:100 |
| Guinea pig anti-PRR19 | a gift from Attila Tóth | N/A | IF | 1:200 |
| Rabbit anti-MSH4 | a gift from Paula Cohen | N/A | IF | 1:200 |
| Guinea pig anti-TEX11 | a gift from Christer Höög | N/A | IF | 1:200 |
| Rabbit anti-MER3 | Rao et al., 2017 | N/A | IF | 1:200 |
| Rabbit anti-RPA2 | Abcam | ab76420 | IF | 1:200 |
| Rabbit anti-DMC1 | Santa Cruz Biotechnology | sc-22768 | IF | 1:50 |
| Mouse anti-RAD51 | ThermoScientific | MS-988-P0 | IF | 1:50 |
| Guinea pig anti-H1t | a gift from Marry A. Handel | N/A | IF | 1:1,000 |
| Human anti-centromere | ImmunoVision | HCT-0100 | IF | 1:500 |
| Mouse anti-p63 | Santa Cruz Biotechnology | sc-25268 | IF | 1:500 |
| Goat anti-mouse 488 | Invitrogen | A-11029 | IF | 1:1,000 |
| Goat anti-rabbit 488 | Invitrogen | A-11070 | IF | 1:1,000 |

| | | | | |
|---|---|---|---|---|
| Goat anti-mouse 594 | Invitrogen | A-11020 | IF | 1:2,000 |
| Goat anti-rabbit 568 | Invitrogen | A-11036 | IF | 1:1,000 |
| Goat anti-human 555 | Invitrogen | A-21433 | IF | 1:1,000 |
| Goat anti-mouse 350 | Invitrogen | A-21049 | IF | 1:50 |
| Donkey anti-guinea pig 488 | JacksonImmunoResearch | 706-545-148 | IF | 1:100 |
| Donkey anti-goat 488 | JacksonImmunoResearch | 705-545-147 | IF | 1:100 |
| Donkey anti-goat Cy3 | JacksonImmunoResearch | 705-165-147 | IF | 1:100 |
| Donkey anti-rabbit Cy3 | JacksonImmunoResearch | 711-165-152 | IF | 1:100 |
| Donkey anti-rabbit 594 | JacksonImmunoResearch | 711-585-152 | IF | 1:500 |
| Donkey anti-mouse 594 | JacksonImmunoResearch | 715-585-151 | IF | 1:100 |
| Donkey anti-rabbit 647 | JacksonImmunoResearch | 711-605-152 | IF | 1:100 |
| Donkey anti-guinea pig 647 | JacksonImmunoResearch | 706-605-148 | IF | 1:100 |
| Donkey anti-goat 647 | JacksonImmunoResearch | 705-605-147 | IF | 1:100 |
| Donkey anti-mouse AMCA | JacksonImmunoResearch | 715-155-151 | IF | 1:50 |
| Donkey anti-rabbit AMCA | JacksonImmunoResearch | 711-155-152 | IF | 1:50 |
| Donkey anti-goat AMCA | JacksonImmunoResearch | 705-155-147 | IF | 1:50 |
| Goat anti-guinea pig IgG(H+L)-HRP | Southern Biotech | 6090-05 | IB | 1:1,000 |
| Rabbit anti-goat IgG(H+L)-HRP | Southern Biotech | 6020-05 | IB | 1:1.000 |

IF, immunofluorescence; IB, immunoblotting.
